# Reproducible processing of TCGA regulatory networks

**DOI:** 10.1101/2024.11.05.622163

**Authors:** Viola Fanfani, Katherine H. Shutta, Panagiotis Mandros, Jonas Fischer, Enakshi Saha, Soel Micheletti, Chen Chen, Marouen Ben Guebila, Camila M. Lopes-Ramos, John Quackenbush

## Abstract

**Background:** Technological advances in sequencing and computation have allowed deep exploration of the molecular basis of diseases. Biological networks have proven to be a useful framework for interrogating omics data and modeling regulatory gene and protein interactions. Large collaborative projects, such as The Cancer Genome Atlas (TCGA), have provided a rich resource for building and validating new computational methods resulting in a plethora of open-source software for downloading, pre-processing, and analyzing those data. However, for an end-to-end analysis of regulatory networks a coherent and reusable workflow is essential to integrate all relevant packages into a robust pipeline.

**Findings:** We developed tcga-data-nf, a Nextflow workflow that allows users to reproducibly infer regulatory networks from the thousands of samples in TCGA using a single command. The workflow can be divided into three main steps: multi-omics data, such as RNA-seq and methylation, are downloaded, preprocessed, and lastly used to infer regulatory network models with the netZoo software tools. The workflow is powered by the NetworkDataCompanion R package, a standalone collection of functions for managing, mapping, and filtering TCGA data. Here we show how the pipeline can be used to study the differences between colon cancer subtypes that could be explained by epigenetic mechanisms. Lastly, we provide pre-generated networks for the 10 most common cancer types that can be readily accessed.

**Conclusions:** tcga-data-nf is a complete yet flexible and extensible framework that enables the reproducible inference and analysis of cancer regulatory networks, bridging a gap in the current universe of software tools.

## Background

There is a growing recognition of the importance of ensuring that scientific research is reproducible and that analytical steps are transparent [1, 2]. The field of bioinformatics has been particularly receptive to this trend, with many prominent scientists and journals advocating for the use of open-source software, open data, and reproducible methods [3, 4]. Projects such as Bioconductor [5] and Bioconda [6] facilitate sharing and reusing bioinformatics software, the Galaxy project [7] has pioneered the development of platforms for training and sharing best practices for complete data analysis workflows, and workflow management tools such as Nextflow [8], Snakemake [9], and WDL [10] facilitate reproducibility of complex data analysis pipelines.

These methodological advances are intertwined with the increasing availability of large-scale biological data. Indeed, the falling cost of sequencing has enabled the generation of population-level omics data, such that thousands of subjects can be profiled in a single project to investigate complex traits and diseases. Notably, the UKBiobank [11] has collected multi-omics and clinical data for more than 500, 000 individuals representative of the UK population, and the 1000 genomes project [12] includes samples from more than 4000 individuals collected to characterize human genetic variation. Ongoing data collection efforts, such as the 100, 000 genome project [13], demonstrates that this data deluge is not slowing down, emphasizing the need for reproducible and strictly managed software pipelines.

The Cancer Genome Atlas (TCGA) [14] was one of the first large collaborative projects designed to study the molecular basis of disease and includes samples collected from more than 10, 000 cancer patients representing more than 30 tumor types. The data from TCGA has been invaluable for the study of regulation in both healthy and tumor tissues [15, 16, 17, 18, 19] and for developing and benchmarking analytical methods for omics data [20, 21]. The TCGA dataset has grown in value with data from related projects such as TCPA [22, 23] and CPTAC [24, 25] creating new opportunities for innovative methods development and applications.

Given that most pathologies are the result of the complex interplay between multiple genomic, transcriptomic, epigenomic and other factors [26, 27], inference and analysis of network models is possibly the most important analytical trend enabled by large-scale omic data resources. Biological networks represent interactions between biological entities to model high-level organization of biological systems and they aim at describing the molecular mechanisms that define biological states and the progression between them. Most notably, network analyses have helped to elucidate the etiology and progression of tumors and provide insight into important features of their clinical manifestation[28, 29, 30, 31, 32]. Many network types have contributed to our understanding of biological processes, including protein-protein interaction networks, networks of DNA-protein interactions, and co-expression networks. Among them, Gene regulatory networks (GRNs) consisting of transcription factors (TFs) and the genes they target for regulation in a particular phenotype have emerged as a particularly powerful tool to describe the complex machinery driving disease development, progression, and response to therapy [33, 34, 35, 36, 37, 38]. Expanding on the idea of characterizing regulation by modeling the interactions between TFs and genes, multi-omics association networks combine multi-modal data to shed light on other factors that can influence regulation. For example, combining transcriptomics and methylomics data [39] can provide insight into whether observed gene expression patterns are linked to DNA methylation, potentially controlled by a proximal or distal element, or whether expression is due to other, potential associations. For these and other applications, the TCGA data is uniquely positioned to be analyzed using network methods that include generating single and multi-omics networks, gene regulatory network models, and by analyzing the relationship between somatic aberrations and their effects on gene regulation.

However, accessing and analyzing, from raw data to networks, the wealth of data that TCGA or similar projects provide is a non-trivial task. For example, raw sequencing data cannot be publicly released and it needs to be aligned and quantified before being used for downstream analyses. Multiomic data require matching samples from different assays, and data pre-processing and filtering steps are fundamental to remove samples that are low quality or population outliers and to identify and remove batch effects. The Genomic Data Commons (GDC) [40, 41] and TCGAbiolinks [42, 43] provide not only programmable access to the data, but also tools to map identifiers and to filter data based on clinical features. For example, with a few lines of code one could download all RNA-seq and mutation data for a specific tumor (or subtype) from those subjects over 70 years of age. The TCGAbiolinks package also provides a number of functions for *ad-hoc* pre-processing of the data, on top of the most commonly used analytical steps such as differential gene expression analysis [44, 43]. Once the data are downloaded and pre-processed, users generally perform additional analyses to extract biologically meaningful insights. The netZoo project [45] actively maintains a growing suite of reusable tools for the inference and analysis of biological networks. Among the sixteen methods currently in netZoo are PANDA [33], which uses gene expression data together with prior TF-binding information and TF-TF interaction data to infers Gene Regulatory Networks (GRNs) describing TF-gene interactions, DRAGON [39], that creates robust multi-omic partial-correlation networks using, for example, matched expression and methylation data, and LIONESS [46] extracts individual networks for each sample in a population using a leave-one-out strategy with linear interpolation.

It is then easy to see that creating an entire analytical pipeline involves combining multiple processing steps. Although there are reusable and documented pieces of software for each specific step, chaining these processes back to back, all the way from data download to final analysis, results in a complex workflow with many, interdependent decisions that must be made as well as tools to facilitate transfer of data and results from one step to the next. To ensure reproducibility, a single robust and transparent workflow is necessary.

We developed *tcga-data-nf* (https://github.com/QuackenbushLab/tcga-data-nf), a Nextflow workflow to generate GRNs with a single command that manages all the steps from data download through pre-processing to network generation. *tcga-data-nf* can download TCGA clinical and phenotypic data, and multi-omic data that include RNA-seq, mutation, methylation and copy number variation data. It then pre-processes these, and generates individual sample GRNs and expression-methylation association networks by combining LIONESS with PANDA and DRAGON, respectively. In a more detailed example (https://github.com/QuackenbushLab/tcga-data-supplement), we show that *tcga-data-nf* not only allows us to swiftly generate networks for the four consensus subtypes of colon cancer [47], but, by combining insights from the various analyses, expand our understanding of the regulatory processes driving prognosis in the more aggressive CMS4 subtype.

As part of developing *tcga-data-nf*, we created the *NetworkDataCompanion* (*NDC*, https://github.com/QuackenbushLab/NetworkDataCompanion) R package that streamlines routine steps in TCGA data processing, including filtering and mapping gene and sample identifiers between modalities (which is often a challenge with such heterogeneous data) and allows modality-specific data transformation, such as normalization and cleaning. *NDC* enables all the pre-processing steps in *tcga-data-nf*, but it is also available as a standalone tool for separate use.

Finally, it is worth mentioning that *tcga-data-nf* comes equipped with all essential supplementary components, including a docker container, conda environments, comprehensive documentation, and introductory tutorials to help users get started. Also, we have uploaded the GRNs for the ten most frequent cancer types (BRCA, LUAD, LUSC, KIRC, LIHC, PAAD, PRAD, SKCM, STAD, COAD) on GRAND [48] such that anyone could access and explore them.

### Findings

The open-source *tcga-data-nf* is a Nextflow [8] workflow that allows users to fully manage gene regulatory network analysis on TCGA data, from the data download and preparation to network inference. By chaining and combining atomic tasks, called processes, this pipeline enables users to infer GRNs and other networks with a single command, only specifying the tumors of interest and the appropriate set of parameters.

The whole workflow can be subdivided in three main functions: downloading the raw data, preparing the data for the analysis, and inferring the networks, see Figure 1. **Download** Data downloaded include RNA-seq, mutation, methylation, and copy number variation data, which are the modalities used to generate regulatory networks and to characterize the DNA aberrations that could explain changes in regulation. Clinical and phenotypic data are also downloaded to support downstream investigations. **Prepare** Gene expression and methylation data are cleaned and pre-processed. Sample duplicates, outliers, and lowly-expressed genes are removed. CpG-level methylation are mapped to overall gene promoter methylation values, and sample identifiers are matched to facilitate integration between different data modalities. **Analyze** The *tcga-data-nf* pipeline enables users to generate tumor-specific GRNs with PANDA [33] and expression-methylation multi-omic partial correlation networks with DRAGON [39]. For both methods, *tcga-data-nf* also facilitates the generation of sample-specific networks with LIONESS [46].

**Figure 1.**
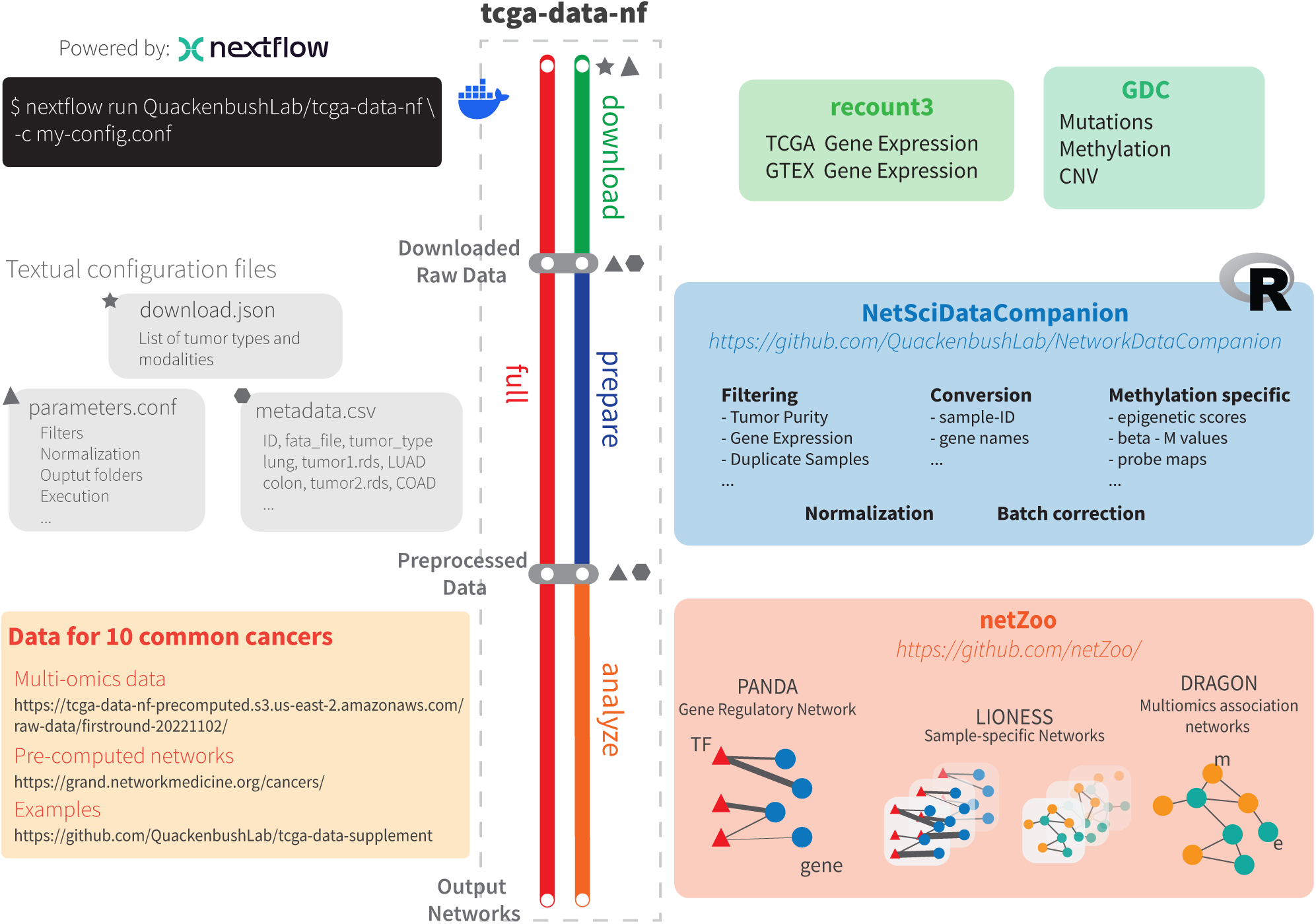
Graphical Abstract

Although designed as a full pipeline that runs all the steps above, *tcga-data-nf* can be used as a modular tool that also allows users to run the download, prepare, and analyze steps independently or in various combinations (Figure 1). The decoupling of different steps is useful from a practical standpoint; the raw data download does not require efficient computational resources, but it is time-consuming and is performed once to generate a long-term data repository, conversely the network inference step generally requires larger amounts of memory and could be run multiple times, as tools evolve, to investigate how various parameters affect the results.

The associated *NetworkDataCompanion* (*NDC*) R package is the engine behind the “Prepare” step of *tcga-data-nf*. While many existing Bioconductor packages implement functions to filter, map, and process TCGA data, it is often necessary to resort to multiple tools to carry out simple functions. The *NDC* package solves this issue by integrating a set of tools into one package, streamlining the routine steps of TCGA data processing. TCGA-specific functions allow users to filter and map gene and sample identifiers between modalities, addressing one often vexing challenge with heterogeneous data. Omic-specific functions allow users to normalize, transform, and filter the data, based on the specific needs of the downstream tasks. Although *NDC* is designed specifically to support *tcga-data-nf*, the package is standalone and can be and reused in other contexts.

State-of-the-art environment and containerization tools are also integrated with *tcga-data-nf*, which can interface with Docker [49], singularity [50], or conda [51]. We provide a general container, (vfanfani/tcga-data-nf) and configuration details for both Docker and Singularity; we also provide a customizable configuration structure, such that all processes can be run with the appropriate conda environments.

Finally, in developing *tcga-data-nf* and *NetworkDataCompanion*, we generated PANDA and PANDA-LIONESS networks for the ten most common tumors in TCGA. Although the analysis we present here uses the colon cancer networks, we provide access to networks for all ten tumor types so they can be used by others without the need to run the more expensive steps of the workflow.

In the following sections we will explain each step of the workflow in more detail, including descriptions of the *tcga-data-nf* pipeline steps and *NetworkDataCompanion* functionalities.

### Download

For the generation and analysis of GRN and association networks, we focus on five data modalities: gene expression, methylation, copy number variation (CNV), and mutation data, alongside patient clinical data. All TCGA data are downloaded from the Genomic Data Commons (GDC project [52]), and we use the “TCGAbiolinks” [43] and “GenomicDataCommons” [40] R packages to manage the download step for methylation, CNV, mutation, and clinical data. However, for the gene expression data, we used recount3 [53] since we were also interested in comparing the results with the GTEx [54] project (which provides “normal” tissue-specific gene expression), taking advantage of recount3’s standardized processing across studies.

Given the variety of data types and the large number of parameters that need to be specified, the download step is driven by a json configuration file (Listing 1). The structure of this file is modality-centric, which means that for each data type (gene expression, mutations, CNVs, …) one can specify the cancer types to be downloaded. The configuration also provides users with the ability to pass a list of samples that one wants to select from the entire population, which is useful to discard problematic samples or focus on a specific subpopulation.

The download step generates metadata tables that contain information about all the downloaded data. These are simple comma-separated tables that store the key parameters used to download each dataset and the path of the resulting files. These tables can be directly used for the following prepare step. A simple scheme of the download step is shown in Supplementary Figure S1.

For this pipeline we have a dedicated testing profile (testDownload) described in the “Testing” section that allows the workflow to be piloted and validated before committing to large and expensive analyses.

### Prepare

The downloaded data needs to be pre-processed before being used in downstream analyses. The array of possible parameter choices regarding normalization and filtering leads to an large number of parameter configurations. Since they can affect the results and conclusions from any analysis[55, 56], we standardized these pre-processing steps and implemented them into the “Prepare” pipeline. The prepare step primarily deals with pre-processing transcriptomics and methylation data, which are those we use most commonly for the generation of GRN and association networks.

### Expression

From recount3, we obtain gene-level raw count data for RNA-seq data from both TCGA and GTEx. In order to clean these data, we implemented the following steps:

- Normalization: raw count data are normalized and one can generate either TPM [57, 58] or CPM (CPM with TMM normalized library size [59, 60]),
- Duplicates: where duplicate samples are present (two or more samples from the same research subject), the sample with the lowest sequencing depth is discarded.
- Batch correction: if specified, batch effect is removed using ComBat [61, 62]. We also visualize the effect of batch removal using PCA.
- Low expression: We remove genes that have low expression, defined as those genes with less than *n* counts in at least *p*% of samples for user-defined values of *n* and *p*.
- Sample purity: for TCGA tumor samples, we remove those that have low purity, using previously computed purity values [63].
- Tissue separation: TCGA contains both tumor and normal samples, and one can save the data for these tissue types separately to allow tumor/normal comparisons.

All these choices are set by appropriate parameters in the configuration file and users can specify multiple values for each parameter so that *tcga-data-nf* outputs results for all parameter combinations.

### Methylation

The TCGA methylation array data undergo two key pre-processing steps: mapping of individual CpG probes to generate gene-level promoter methylation values and general data cleaning/transformation. By default, CpGs are mapped to genes using the publicly available annotation for the Illumina 450k array, mapped to hg38 using Gencode v36 (https://zwdzwd.github.io/InfiniumAnnotation) although users can supply their own annotation file if desired. Once CpGs are mapped to genes, the average promoter methylation for each gene is calculated. The promoter region is defined as the area 200 basepairs upstream of the transcription start site (users can redefine this boundary), and average of the methylation beta values for any probes falling within this region is calculated.

It is not necessary to perform this mapping to promoter methylation over all the genes represented on the EPIC array but rather to a particular subset that may be relevant for a particular analysis. For example, in the application of DRAGON [39] that we describe, we map the probes only to genes encoding transcription factors (TFs).

After obtaining gene-level methylation for each sample, we preprocess the data as follows:

- Duplicates: If a sample has multiple methylation array profiles, we select one of these at random for downstream use and discard the remainder.
- Missing data: Any gene that has missing promoter methylation values for more than 20% of the samples is removed from the analysis.

If a gene has missing values for ≤ 20% of the samples, these missing values are estimated by mean imputation.

- Conversion from beta to M-values: The mean promoter methylation beta value β ∈ [0, 1] is converted into an M-value *M* ∈ (–∞, ∞) using the formula:

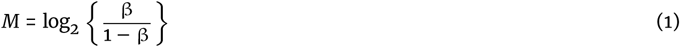

See [64] for a detailed discussion of the relative merits of using β-values vs. M-values in methylation analyses.

- Transformation to approximate normality: We apply a nonparanormal transformation [65] to the M-values to achieve approximate normality in the distribution for input into DRAGON as it requires approximately normal data. The software used for this transformation is the huge.npn function of the R package huge [66].

A simple scheme of the prepare step is shown in Supplementary Figure S2. At the end of these steps, we obtain a comma-separated file, with rows being the probes and columns the samples. These processing steps rely on the companion R package *NetworkDataCompanion* (*NDC*) that is described in depth in the “*NetworkDataCompanion* “” section.

### Analyze

Finally, once transcriptomic and methylation data are pre-processed, we can generate the regulatory networks. The analyze pipeline generates both GRNs with PANDA and multi-omic association networks with DRAGON. PANDA [33] is a GRN inference method that uses bulk expression data, together with a prior TF-gene binding network (based on TF motif mapping) and a TF-TF protein interaction network, and generates population-level bipartite TF-gene regulatory networks. DRAGON [39] is a partial correlation network inference method based on Gaussian graphical models that infers multi-omic associations between different omics modalities. We also used LIONESS [46], which estimates sample-specific networks by interpolation between a network for the entire population and that for the population less the sample for which we are estimating a network. LIONESS is agnostic to network estimation; the choice of the method is delegated to users who can specify the method in the configuration file. In our pipeline, we used LIONESS to estimate both PANDA and DRAGON networks for each sample. A simple scheme of the prepare step is shown in Supplementary Figure S2.

The *analyze* workflow relies on implementations of PANDA, DRAGON, and LIONESS in the netZooPy package [45]. The processes for network inference uses the command line interfaces and Python objects for PANDA, DRAGON, and LIONESS; these require users to specify the expression/methylation input files and the parameters for each method. Conveniently, the prepare step in *tcga-data-nf* generates a metadata table that the users can directly employ to specify which files are then used by the analyze step. Given the amount of data the pipeline can generate, we also updated netZooPy to save networks in Hierarchical Data Format (HDF) which reduces the size and reading-writing time for storing the networks.

Lastly, we note that although we have compiled a pipeline of methods that use gene expression and methylation to generate network models, the workflow can be easily edited to include other data types and methods.

### Full

The full pipeline combines the download, prepare, and analyze steps described above. It is designed to be run with a single command, and it represents most complete network analysis pipeline. While we recommend separating the three steps, for instance by downloading the data once and then running the prepare and analyze steps as needs change, there are instances where researchers might want to gather and analyze all the data for a single project at once. In section, we show an example of how the full pipeline can be used to generate DRAGON and PANDA networks for the TCGA colon cancer consensus subtypes.

A simple scheme of the full steps is shown in Supplementary Figure S4. For the full pipeline, the configuration files and parameters mirror those of the three modular pipelines above. First, a “json” configuration file, see Listing 2, similar to the one for downloading data, needs to be populated with all the data modalities of interest. Then, the processing and analysis parameters need to be specified in the nextflow configuration file.

### NetworkDataCompanion

The *NetworkDataCompanion* (*NDC*) R package supports pre-processing of TCGA bulk RNA-seq and DNA methylation data. This version-controlled R package for these functions helps us attain the high standard of reproducibility in the overall *tcga-data-nf* pipeline. While *NDC* is the engine behind the *tcga-data-nf* workflow processing steps, the software is standalone and can be installed and used outside the workflow. *NDC* currently provides three broad classes of functions: functions for mapping identifiers, functions for filtering data, and functions for preparing expression and methylation data (such as normalization and scaling) (Figure 1). While many of the functions described below are intuitively simple, it is worth noting that the TCGA project, with its wealth of data, requires checks on sample quality and multi-omic sample mappings.

### Mapping functions

Given the variety of data types used in the pipeline and the number of other resources with which they interface, there are many instances where we need to map between “synonymous” identifiers. We have implemented several wrapper functions that use existing tools such as GDC and TCGAutils to retrieve and convert various sample identifiers (TCGA barcodes, UUIDs). We have also implemented functions for translating gene names between Ensemble IDs [67], HUGO Gene Nomenclature [68], and Entrez IDs [69, 70] by leveraging Gencode v26 [71] which was used for TCGA and the AnnotationDbi R package [72].

#### Sample filtering functions

By default, *tcga-data-nf* downloads and processes all samples available for a particular TCGA dataset. *NDC* provides three different functions that allow a user to filter these samples. First, a user may eliminate duplicate samples using a method that selects the sample with the strongest signal based on RNA sequencing depth. Second, a user may filter samples based on the TCGA sample type (for example, primary tumor tissue, metastatic tissue, or adjacent normal tissue). Finally, a user may filter samples based on tumor purity, excluding samples where the number of non-tumor cells is too large according to sample published annotation [63].

### Data preparation functions

Lastly, with *NDC* users can apply common data transformations to both gene expression and methylation data. For RNA-seq data from recount3, one can normalize read counts to transcripts per million (TPM) [57] or counts per million (CPM) [60] and their corresponding log transformations. For methylation data, functions are provided to convert methylation beta values to m-values (logit base-2 transformed beta values) and vice-versa [73] and to aggregate methylation values to get gene-level information, such as average methylation within a promoter region or gene body.

Collectively, the functions in *NetworkDataCompanion* represent a comprehensive set of tools for basic processing, filtering, and mapping that are needed to clean and prepare TCGA data. We note that although there are R packages that cover each of the individual tasks described above, *NetworkDataCompanion* aggregates them into a unified solution in one version-controlled package to facilitate their use together and to allow them to be seamlessly integrated into the *tcga-data-nf* pipeline.

### Multiomics associations identify changes between colon subtypes

Colorectal cancer affects nearly two million individuals worldwide each year and is forecast to increase in incidence by 84% in the next 20 years [74]. Molecular profiling studies have identified four major expression-based consensus subtypes (CMS1, MSI immune, CMS2, canonical, CMS3, metabolic, and CMS4, mesenchymal) [47] with distinct phenotypic and clinical features. CMS1 and CMS3, each of which has a distinctive genomic and epigenomic profile, together represent around 25% of cases. CMS2 and CMS4 are the most common subtypes, and although they have similar patterns of somatic mutation, structural variation, and methylation, they differ significantly in outcomes; CMS2 tumors have good survival rates, while CMS4 cases are characterized by more aggressive tumors and poorer prognosis [75]. Additional studies have described intra-tumor heterogeneity and clinically actionable features that distinguish CMS4 and have found evidence that this more aggressive mesenchymal subtype often arises from CMS2-like tumors [76]. We then reasoned that multiomics association networks and GRNs could provide further insight into the mechanisms that drive the difference between these subtypes.

Using the full pipeline, we specified the TCGA samples that belong to each subtype [47] and were able to download and pre-process the data and generate the networks for each of them. We began with the four subtype-specific DRAGON partial correlation networks linking TF expression to promoter methylation. Given *F* transcription factors for which we have both methylation and expression data, each network has 2*F* nodes, is symmetric, and can be divided into three blocks: i) expression-expression correlations, homogeneous (*E_i_*, *E_j_*) edges between genes *i* and *j*, ii) methylation-expression correlations, heterogeneous (*M_i_*, *E_j_*) edges between genes *i* and *j*, and iii) methylation-methylation correlations, homogeneous (*M_i_*, *M_j_*) edges between genes *i* and *j*.

Because methylation is generally inhibitory, we expect promoter methylation to be inversely correlated with gene expression [77, 78]. Indeed, by examining the distribution of methylation to expression edge weights (*M_i_*, *E_j_*), we see that weights for edges connecting a gene to the methylation status of its promoter (*M_i_*, *E_i_*) have a distribution more strongly skewed to negative values than is the distribution connecting one gene to the methylated promoters of other genes (Figure 2A). One can also see that there is good correlation between the values of these “same-same” (*M_i_*, *E_i_*) edges between different subtypes (Supplementary Figure S5). Some of “same-same” genes (STAT5A, CREB3L1, ZNF24, HMGA1, IRF8, PAX8, CDX2, CREB3L2, TFEB, MGA, NFIB, KLF6, LEF1, HOXD13, HOXA13, HOXB13, GATA2), for which we have evidence of possible epigenetic control, are known to be cancer drivers [79, 80]. For example, STAT5A, the only TF that has low (*M_i_*, *E_i_*) edges in all subtypes, is a known oncogene involved in the JAK signaling cascade [81, 82], while CREB3L1, LEF1, PAX8 are all known to be involved invasion and metastasis [83, 84, 85, 86].

**Figure 2.**
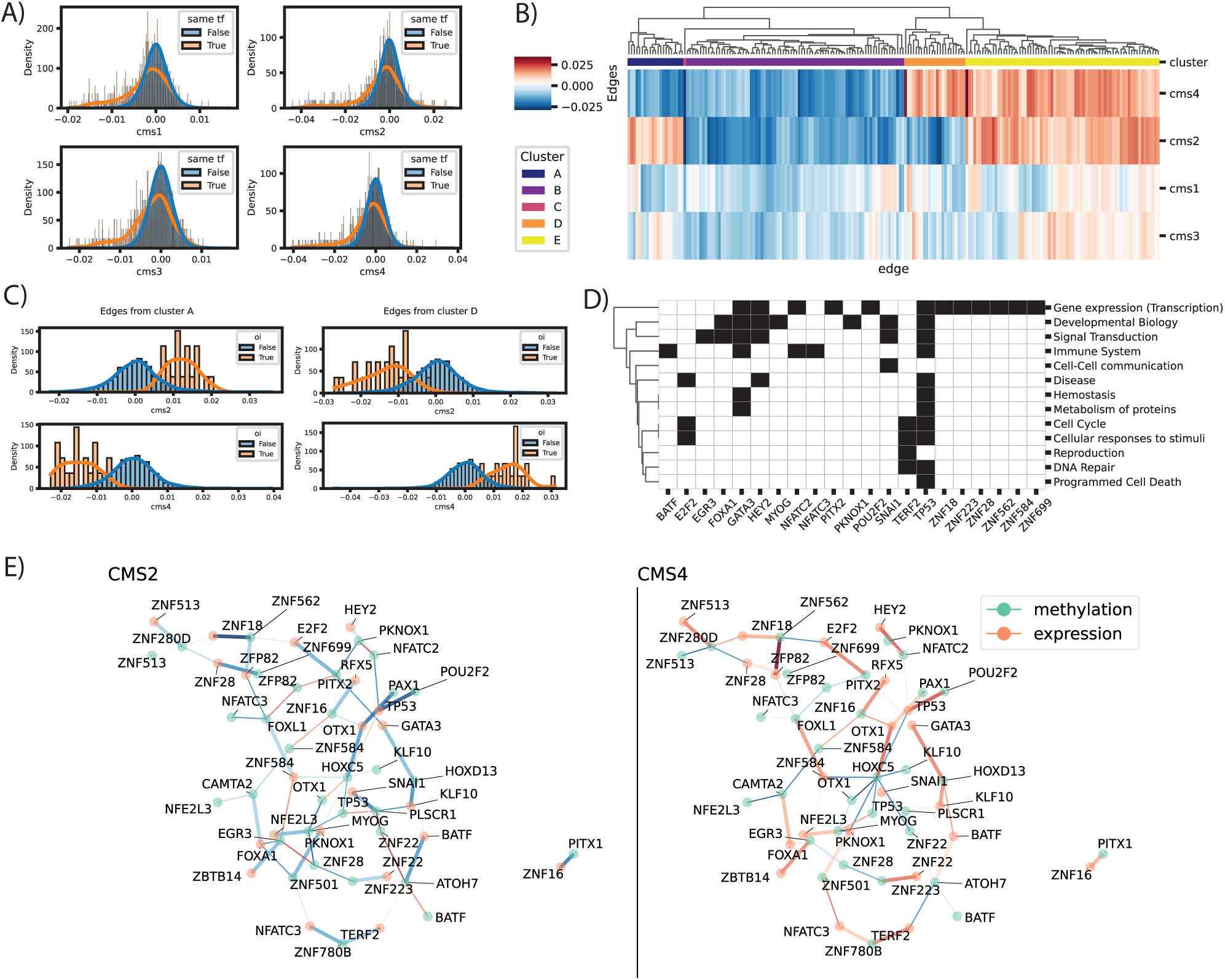
Differences in methylation-expression association of TFs in colon cancer subtypes. A) Distribution of partial correlation values between methylation and expression of TFs in all subtypes. In orange we show the values for the edges of the same TF, that is the correlation between the methylation of the promoter and the expression of the that same TF. As expected, methylation and expression tend to be negatively correlated. The histogram represents the distribution density and it is normalized per subtype and per group. B) For inter-modality edge weights (methylation to expression) we remove the edges between the same transcription factor, and we select the first 200 strongest edges (highest average absolute value of correlation) and we cluster them by correlation values for each subtype (average linkage, Euclidean distance). Interestingly, for cluster A and D the edge values for CMS2 and CMS4 are swapped in direction. C) Distribution of the edge weights for the TFs in clusters A and D. We select first the subgraphs with all the nodes in each cluster, it is worth noting that these graphs contains both the edges represented in figure B, and those that connect the nodes in the cluster but were not the strongest edges. We compare the values of the edges of interest (orange, oi), that are those shown in figure B, and the rest of the edges (blue) that connect the same TFs. We observe that for both groups the average edge value is around 0, while the edges that were selected in the clusters have higher and lower values for CMS2 and CMS4. D) Annotation of all TFs in cluster D (columns) to the Reactome parent term. “Immune system” and “Cellular respondes to stimuli” are more consistenly involved in cluster D, in comparison to cluster A. Here we can discern which TFs are annotated to each term, for instance a a good number of TFs are involved in the immune system (BATF, GATA3, NFATC2, NFATC3, TP53). As expected many of the TFs are annotated to generic transcription pathways, and TP53 is annotated to almost all terms.E) Association graph for the TFs in cluster D for CMS2 (left) and CMS4 (right). We have selected the edges that belong to cluster D (thicker edges) and we have added other 20 top edges, by absolute value, that connect the same TFs (thinner edges), such that we have a connected graph. We show the TFs as nodes with different colors for methylation (green) and expression (orange), and we show the edge values color-coded by the correlation value. Both graphs have the same edges, but it is clear that for many of them, the direction of the correlation is different between the two subtypes.

We then explored the heterogeneous partial correlation edges (*M_i_*, *E_j_*) that connect methylation associated with one TF to the expression of another TF and focused on the edges that differ between the subtypes, under the hypothesis that such edges can identify complex regulatory relationships that distinguish disease states. We selected the edges with the highest absolute value (either the strongest positive or negative correlation) and used hierarchical clustering (average linkage, Euclidean distance) on these edges. We identified five clusters (Figure 2B) with clusters A and D showing distinct patterns that represent reversed associations between CMS2 and CMS4. Even when compared to the rest of the edges connecting the same TFs (Figure 2C), these edges switch direction between the two suggesting a specific change in the regulatory process. Using functional enrichment analysis, we found that both clusters A and D are involved in the metabolism of proteins and DNA repair (Supplementary Figure S6). However cluster D, in contrast to cluster A, includes genes preferentially involved in the Immune System, through the TFs BATF, GATA3, NFATC3, NFATC2 and TP53, as well as cellular responses to stimuli mediated by TFs E2F, TERF2, and TP53 (Figure 2D). This over-representation of immune-related TFs is consistent with reports that CMS4-like tumors exhibit greater degree of immune infiltration than do other subtypes [76]. Further, epigenetic changes in the FOX/HOX and SNAI TF families are known to be involved in colorectal cancer etiology [87], and TP53, FOXA1, GATA3, and GATA6 are all TFs that, when active in an aberrant form, are recognized as being hallmarks of cancer [88]. All these differences between CMS2 and CMS4 can also be visualized at once as the subgraph that emerges from cluster D, Figure 2E.

The DRAGON networks that we inferred here capture information about patterns of DNA methylation and how these are associated with transcription factor expression. The working hypothesis behind that analysis is that some transcription factors exhibit altered patterns of methylation that affect gene expression and ultimately exert downstream effects that help to define different cancer subtypes. Indeed, we identified a number of TFs that differ between the CMS2 and CMS4 subtypes and provide a plausible explanation for the differences they present in the clinic. However, the associations emerging from DRAGON’s partial correlation Gaussian graphical models do not capture the actual regulatory effects that TF have on their target genes.

PANDA is a GRN inference method that uses prior knowledge on motif binding and TF-TF physical interactions together with assayed gene expression to generate a bipartite graph associating TFs with the genes they likely regulate. Each TF-gene edge is weighted by a measure of the evidence of a regulatory relationship. After generating PANDA GRN networks, we performed an analysis directed at identifying the regulatory context for the TFs identified in clusters A and D, those for which we have evidence of changes in methylation and expression patterns between CM2 and CMS4. After having identified the TFs of interest with the DRAGON analysis, we proceeded to investigate the functional role of their targets. We compared the two PANDA networks and selected the edges with the most significant changes between the two subtypes, to identify differentially regulated targets. A gene set enrichment analysis on these differentially targeted genes reveals that they are involved in the transcriptional misregulation of cancer and various immune-related REACTOME pathways that include “Creation of C4 and C2 activators,” “Initial triggering of complement,” and “Signaling by the B Cell Receptor (BCR)” (see Supplementary Tables S1,S2). Moreover, TFs in cluster D seem to consistently differentially target genes in the TGF-*beta* signaling pathway, Cytokine-cytokine receptor interaction and Antigen processing and presentation (Supplementary Figure S7 and Supplementary Tables S3, S4, which is consistent with the observation that TGF-β is not only associated with poorer prognosis in colon cancer, but it is also a involved in the transition between CMS2 to CMS4 [75].

### Testing

Apart from the main workflows for TCGA data analysis, which we showcased in the previous section, our framework also offers a testing option that assesses the general availability and formatting of data and software and runs a trial analysis on a sample dataset.

Nextflow profiles are sets of configuration parameters accessible with the “–profile” scope. For testing purposes, we provide testing configuration profiles for all the workflows with the following names: “test” for the full pipeline, and “testDownload”, “testPrepare”, “testAnalyze” for the specific workflows.

#### test

This profile runs the full *tcga-data-nf* pipeline on a subset of TCGA Pancreatic Adenocarcinoma data (PAAD). All parameters are specified in the (‘conf/test.config‘)

#### testDownload

To test the download step, we need to confirm that the TCGA data are available and retrievable and hence we cannot use minimal dummy datasets. To avoid downloading files that are too large, during testing we only retrieve data from TCGA Pancreas Adenocarcinoma, one of the smallest datasets. All data are downloaded into the ‘results/download_test/‘ folder.

#### testPrepare

To test the prepare step, dummy expression and methylation datasets are provided that contain randomized sample labels and values. Both datasets have only 10 samples each, hence this step should compute quickly. To further simplify this test, each of the parameters that we pass to the recount pipeline is tested for only one value, such that only one input configuration is tested and only one output is produced.

#### testAnalyze

To test the analyze step, we provide dummy preprocessed data for expression and methylation data in the testdata/analyze_ expression.csv and testdata/analyze_methylation.csv files. Alongside the input data, we also provide TF-target binding and PPI network dummy data which are necessary to infer PANDA networks. The test requires that all network methods (PANDA, DRAGON, and LIONESS) are computed in order to confirm that the whole implementation is functioning properly and the netZooPy package is correctly installed and running.

## Discussion

There is a growing recognition that inferring and analyzing gene regulatory network models can provide unique and verifiable insights into the drivers of disease [34]. The GRAND database provides access to freely available, genome-wide gene regulatory network models for samples in TCGA [48], but these models are derived with a fixed set of model parameters and data processing choices that may differ from those that are optimal for a particular application–including choice of normalization method [89] or sample and gene filtering [90]. Further, network inference using tools such as PANDA, DRAGON, or LIONESS are dependent on intermediary files that can be useful for exploratory data analysis and validation tests. For those wishing to generate GRN models using data from TCGA, setting up the requisite environments and data structures can present some challenges, even for those with experience in bioinformatics and computational biology.

In this manuscript, we describe an end-to-end reproducible workflow to download, pre-process, and generate regulatory networks from TCGA cancer data with a single command. This workflow is publicly available, uses fully open-source software, and adheres to what has been deemed the gold standard for reproducibility [3]. The workflow allows the pre-processing of multi-omic data and inference of regulatory networks without requiring users to write code *de novo* and instead only asking them to specify a small number of parameters. We also provide resources in addition to the workflow itself to make implementing the process smoother, including Docker containers and conda environments, extensive configuration files, documentation describing how to reuse the workflow, and pre-generated networks for the ten most common cancer types in TCGA.

As a demonstration of the values of the *tcga-data-nf* pipeline and its flexibility, we showed how multi-omics association networks and GRNs can be used to study transcriptional colon cancer subtypes. We found evidence of previously undescribed methylation-gene expression interactions that target TGF-β signaling and that may help to explain factors influencing the transition between subtypes CMS2 and CMS4.

The biggest hurdle in designing *tcga-data-nf* has been the trade-off between flexibility and completeness. We reasoned that individual pieces of software, such as GDC, TCGABiolinks, edgeR, and netZooPy already provide a broad set of functions covering all the steps required for network generation and analysis. As such, the workflow was designed to chain these tools seamlessly together, allowing users to carry out complex analyses simply by specifying a small number of parameters. While flexible by design, the release version of *tcga-data-nf* does not provide users with unlimited options. We provide only three common gene-expression normalization methods, the full workflow can only be applied to single tumor types and cannot generate pancancer analyses (unless using an *ad-hoc* analyze step), and the workflow does not cover all possible data types available from TCGA—such as miRNA expression. However, we organized *tcga-data-nf* so that it can be easily extended to include other data types and analysis methods and we provide individual download, prepare, and analyze steps that can be used separate from the rest of the pipeline.

## Methods

### Pathway Analysis

In section we carried out all pathways analysis using the GSEApy package [91]. We tested for over-representation (ORA) of TFs and genes in both the Reactome and KEGG sets of pathways with a hypergeometric test. For both cases we selected the appropriate background, that is all TFs in the DRAGON networks or the gene targets in the PANDA networks. The KEGG dataset was downloaded from the GSEApy package as “KEGG2021” dataset.

From the REACTOME database we downloaded the pathway files on June 18th 2024. We have downloaded the tables that map each gene identifier to a pathway, and that also map each pathway to the parent terms. For instance “Intracellular signaling by second messengers“, “Signaling by GPCR“, “Signaling by Hedgehog“… are all part of the “Signaling Transduction” term. To reduce the number of tested pathways, which also reduces overlaps between pathways, we have generated a “slim” set of pathways. For each leaf in the reactome dataset, we have kept only the “parent” node. This way we avoid keeping all the nodes that are too small, and we keep only the depth-1 term. All code and data used for the ORA are in the https://github.com/QuackenbushLab/tcga-data-supplement/ repository.

### Reference data

PANDA uses prior knowledge on putative TF-motif binding and TF-TF contact. To create the regulatory motif network, we downloaded transcription factor motifs for *Homo sapiens* with direct or inferred evidence from the Catalog of Inferred Sequence Binding Preferences (CIS-BP) Build 2.0, accessible at http://cisbp.ccbr.utoronto.ca. These transcription factor position weight matrices (PWM) were mapped to the human genome (hg38) using FIMO [92]. We retained only highly significant matches (*p* ≤ 10^-^5) occurring within the promoter regions of Ensembl genes (specifically, GENCODE v39 annotations retrieved from http://genome.ucsc.edu/cgi-bin/hgTables). These promoter regions were defined as the interval of [-750; +250] base pairs centered around the transcription start site (TSS). This process yielded an initial set of potential regulatory interactions involving 997 transcription factors that collectively targeted 61,485 genes.

For the TF-TF cooperativity prior we obtained PPI data from the StringDB database (version 11.5) using the STRINGdb Bioconductor package [93]. Subsequently, we filtered the PPI data to retain only interactions between transcription factors in the TF-motif network (using a score threshold index of 0). To maintain consistency in PPI scores, we normalized them by dividing each score by 1000, thereby restricting the values to a uniform range of 0 to 1 for both the PPI dataset and the TF-motif network. Additionally, we set self-interactions between transcription factors to a value of one. Since PPI networks are inherently undirected, we transformed the data into a symmetric PPI matrix.

### Availability of source code and requirements

#### tcga-data-nf

- Project name: *tcga-data-nf*
- Project home page: e.g. https://github.com/QuackenbushLab/tcga-data-nf
- Operating system(s): e.g. Platform independent
- Docker: https://hub.docker.com/r/violafanfani/tcga-data-nf
- Programming language: Nextflow, R, Python, bash
- Other requirements: Java, Nextflow
- License: GNU General Public License v3.0

#### NetworkDataCompanion

- Project name: *NetworkDataCompanion*, *NDC*
- Project home page: e.g. https://github.com/QuackenbushLab/NetworkDataCompanion
- Operating system(s): MacOS, Linux
- Programming language: R
- License: GNU General Public License v3.0

#### Notebooks and configuration files

We provide a GitHub repository that contains i) all configuration files mentioned in this manuscript ii) Notebooks and supplementary data for the analysis of colon cancer subtypes.

- Project name: tcga-data-supplement
- Project home page: e.g. https://github.com/QuackenbushLab/tcga-data-supplement
- Operating system(s): Linux, MacOS, Windows
- Programming language: Python
- License: MIT

## Data availability

We precomputed networks for Breast invasive carcinoma (BRCA), Lung adenocarcinoma and Lung squamous cell carcinoma (LUAD, LUSC), Kidney renal clear cell carcinoma (KIRC), Liver hepatocellular carcinoma (LIHC), Pancreatic adenocarcinoma (PAAD), Skin Cutaneous Melanoma (SKCM), Stomach adenocarcinoma (STAD), Colon adenocarcinoma (COAD), and Prostate adenocarcinoma (PRAD). Raw, multimodal data, and processed data are available at https://tcga-data-nf-precomputed.s3.us-east-2.amazonaws.com/raw-data/firstround-20221102 and a guide to download the data is available at https://github.com/QuackenbushLab/tcga-data-supplement/blob/main/data/manifests/manifests.md. PANDA and PANDA-LIONESS networks are are available on GRAND https://grand.networkmedicine.org/cancers/ [48]. Replication data for the subsection are stored on the Harvard Dataverse (https://doi.org/10.7910/DVN/MCSSYJ).

## Declarations

### Funding

This work was supported by grants from the National Institutes of Health: ES, CMLR, MBG, VF, JF, KHS, PM, CC, SM and JQ were supported by R35CA220523; MBG and JQ were also supported by U24CA231846; JQ received additional support from P50CA127003; JQ was supported by R01HG011393; KHS was supported by R01HG125975 and P01HL114501 and T32HL007427; CMLR was supported by K01HL166376; CMLR and ES were also supported by the American Lung Association grant LCD-821824.

### Author’s Contributions

**Conceptualization:** VF, ES, PM, JF, KHS, CMLR and JQ; **Methodology:** VF, KHS, PM, JF, SM, CC; **Software VF, KHS, PM, JF, SM, CC**; **Formal Analysis:** VF; **Resources:** JQ, CMRS, and MBG; **Data Curation:** VF, ES, KHS, PM, and CMLR; **Writing – Original Draft:** VF, KHS; **Writing – Review and Editing:** PM, MBG, JF, KHS, CC, SM, CMLR and JQ; **Visualization:** VF; **Supervision:** JQ, CMLR; **Funding Acquisition:** JQ and CMLR.

## Supplementary

### Download

Listing 1. download-test.json, Example of a json configuration file for the download step. With this file the pipeline downloads all modalities for TCGA LUAD and gtex lung.

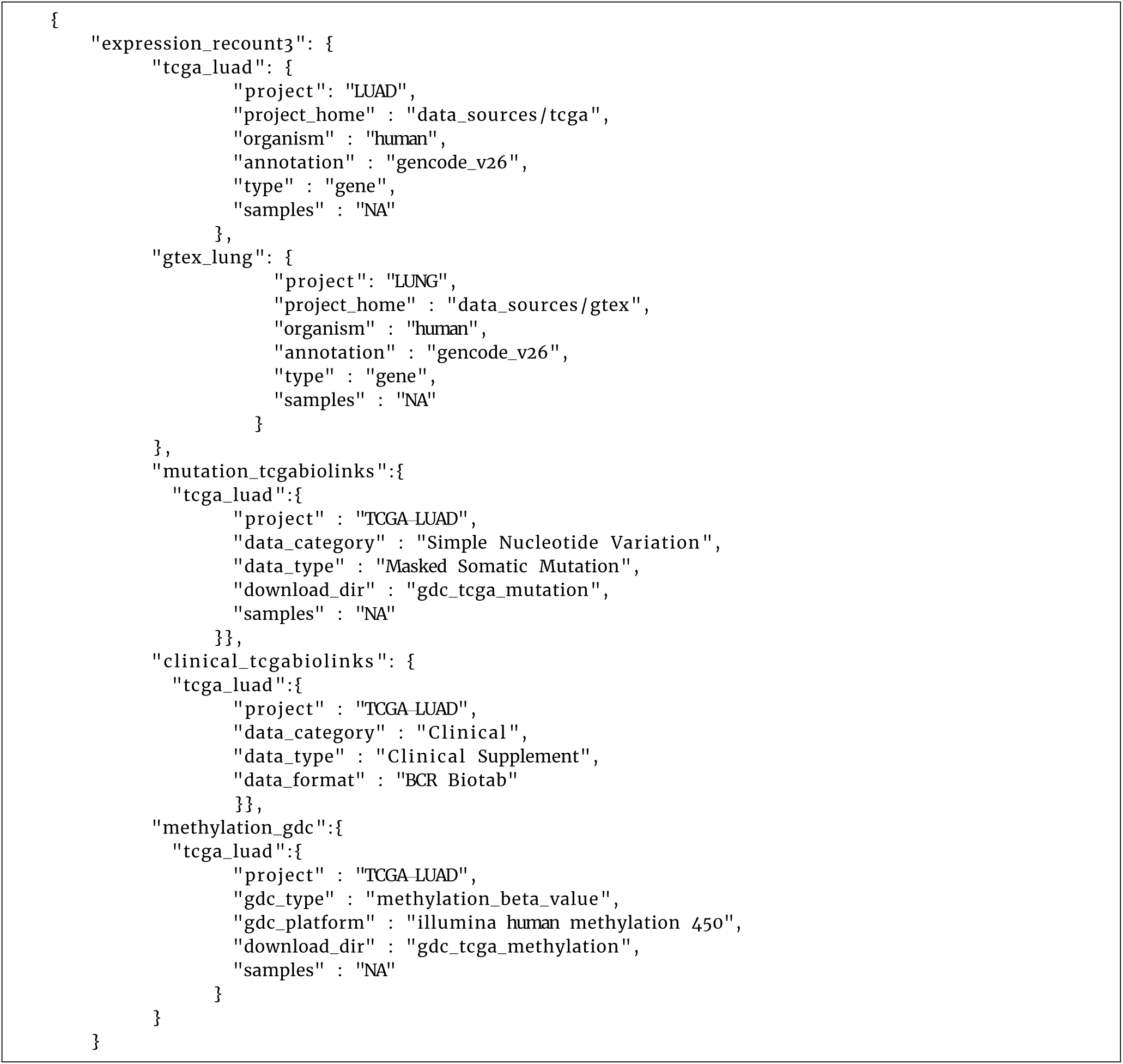

Listing 2. full-test.json, Example of a json configuration file for the full pipeline. With this file we specify which modalities and samples need to be downloaded, pre-processed and analyzed. In this case we are interested in LUAD samples.

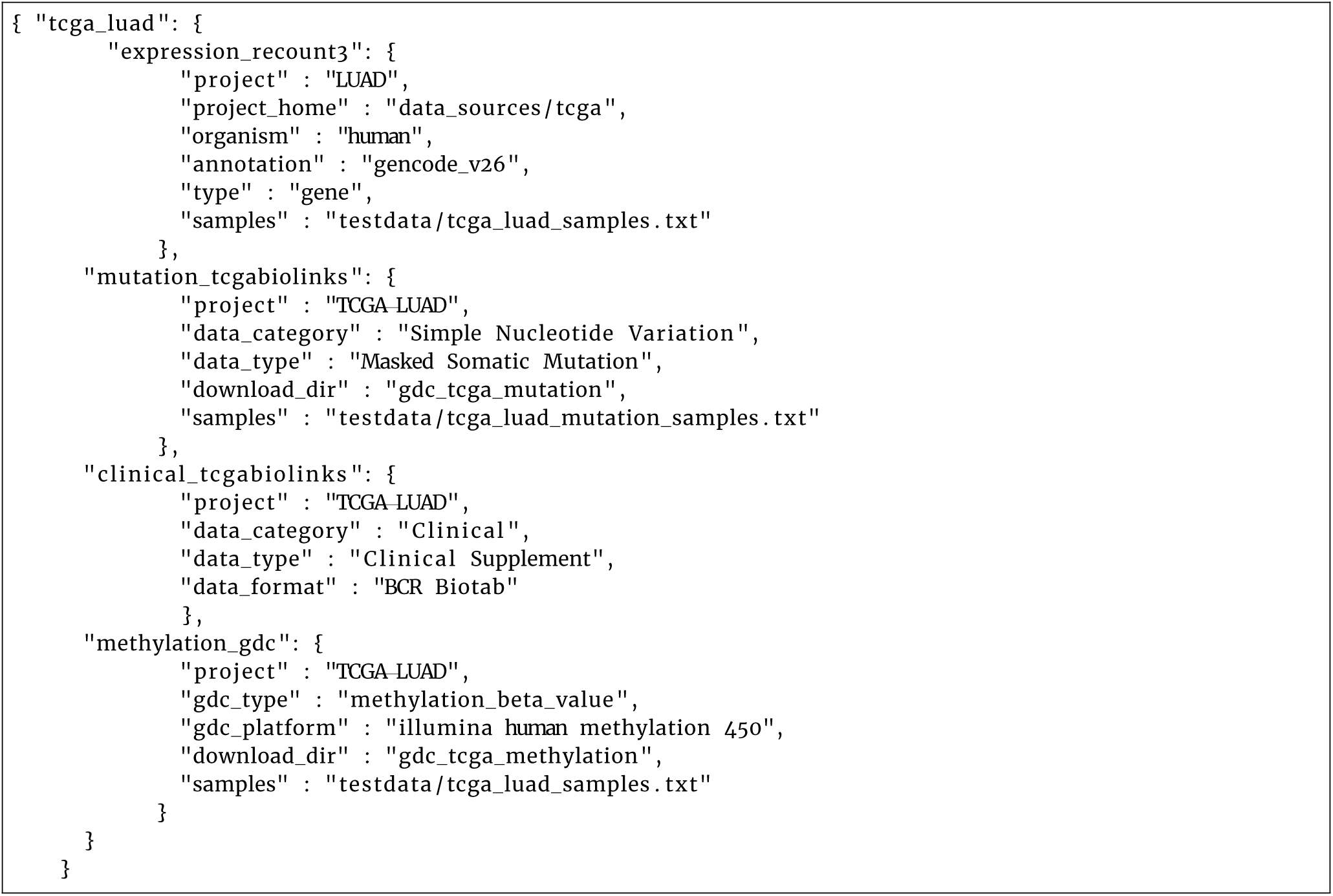

## Supplementary Figures

**Figure S1.**
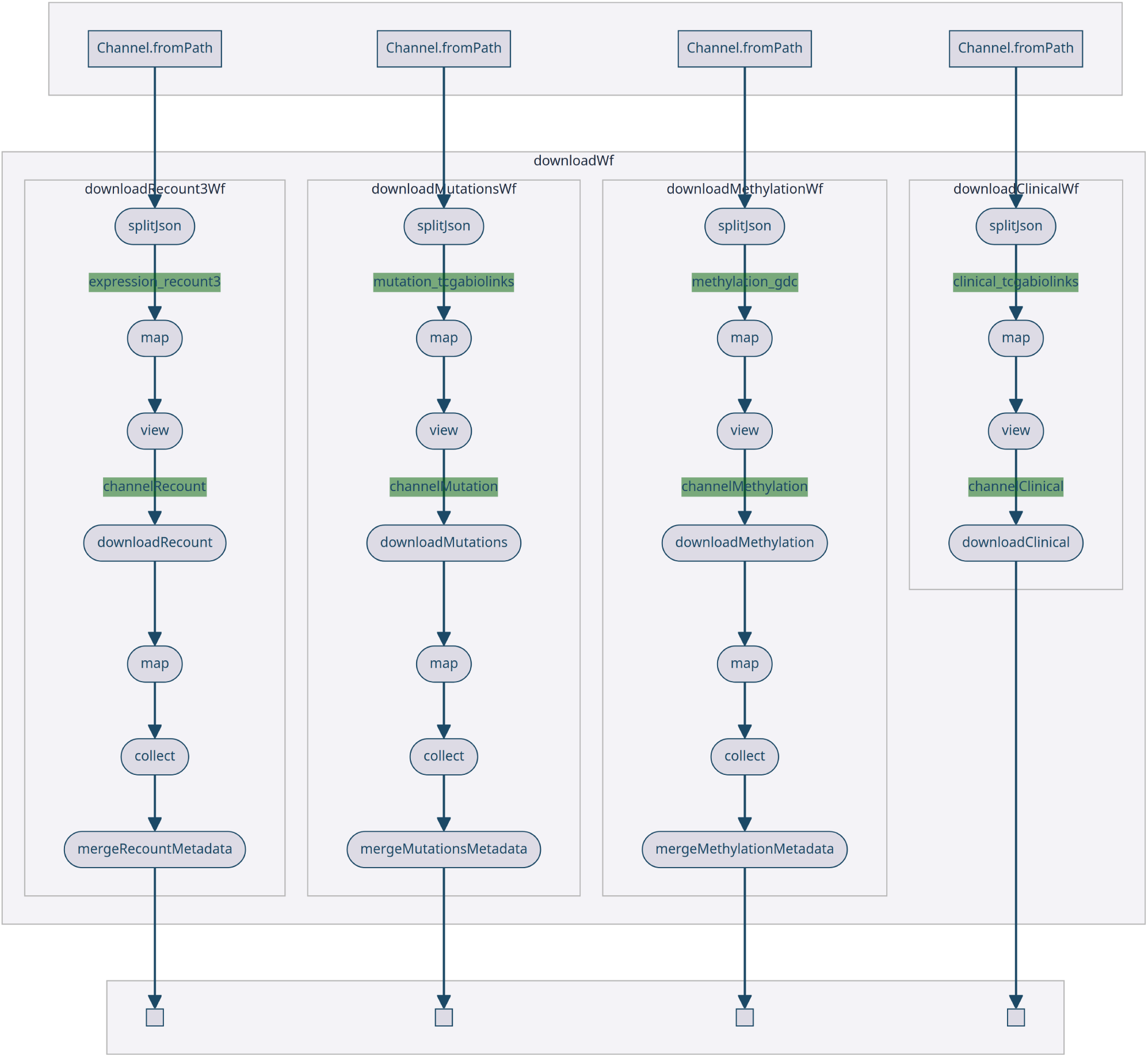
Download. Directed Acyclic Graph of the processes specified in the download pipeline. For each modality that is specified in the configuration file, *tcga-data-nf* downloads the data and generates metadata tables with the names, paths, and parameters of the files. Whenever *tcga-data-nf* is run, we also generate and save the configuration parameters which can be then examined and reused (saveConfig process).

**Figure S2.**
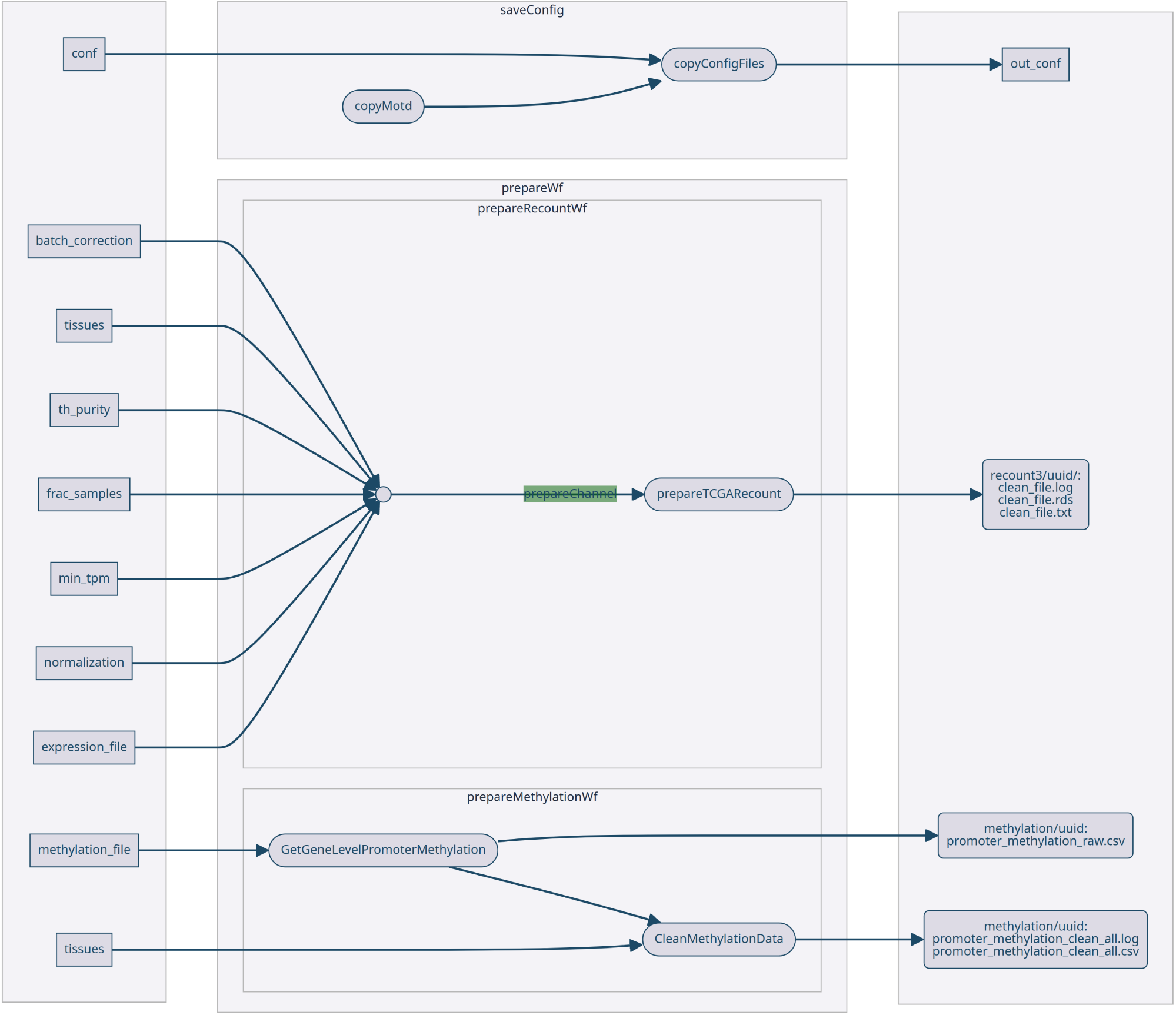
Prepare. Directed Acyclic Graph of the processes specified in the prepare pipeline. The expression and methylation data specified in the configuration metadata is processed using the combination of all input parameters (tissues, purity, minTPM…). Whenever *tcga-data-nf* is run, we also generate and save the configuration parameters which can be then examined and reused (saveConfig process).

**Figure S3.**
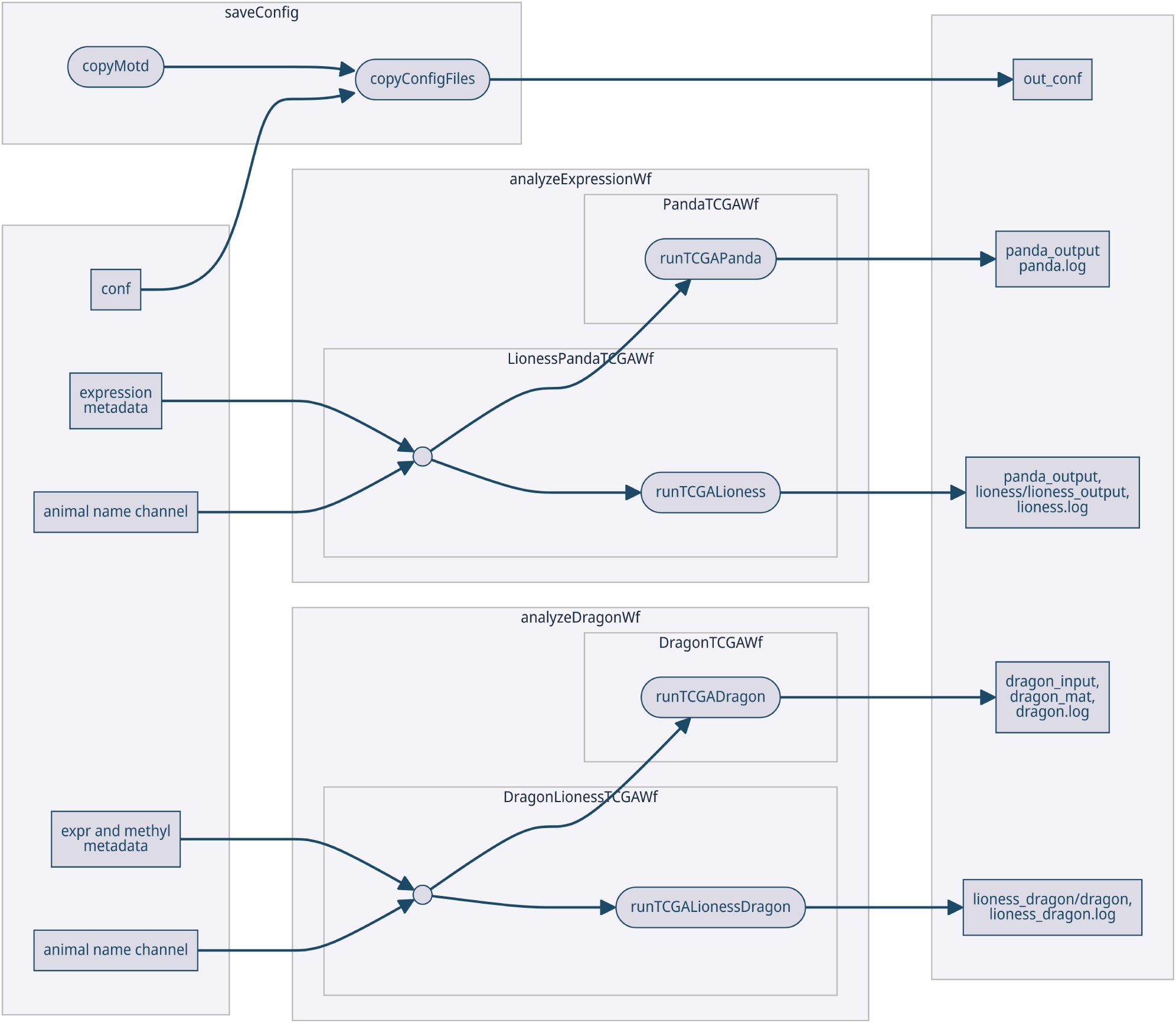
Analyze. Directed Acyclic Graph of the processes specified in the analyze pipeline. Using the input metadata, the *tcga-data-nf* workflow generates PANDA, DRAGON and LIONESS networks, and matches them with log files and intermediate tables, useful for further investigation of the results. Whenever *tcga-data-nf* is run, we also generate and save the configuration parameters which can be then examined and reused (saveConfig process).

**Figure S4.**
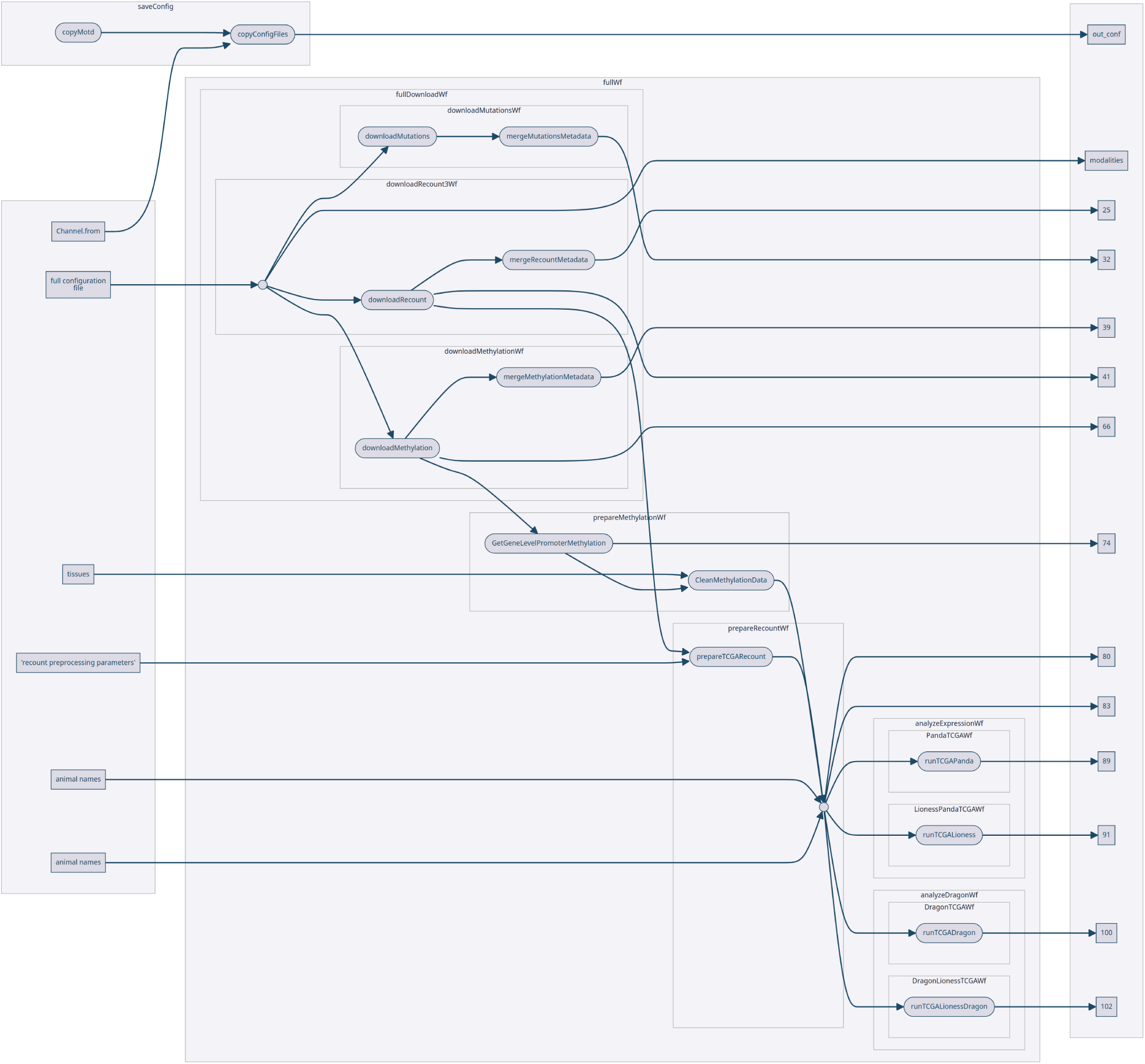
Full. Directed Acyclic Graph of the processes specified in the full pipeline. This workflows combined the download, prepare, analyze steps. Whenever *tcga-data-nf* is run, we also generate and save the configuration parameters which can be then examined and reused (saveConfig process).

**Figure S5.**
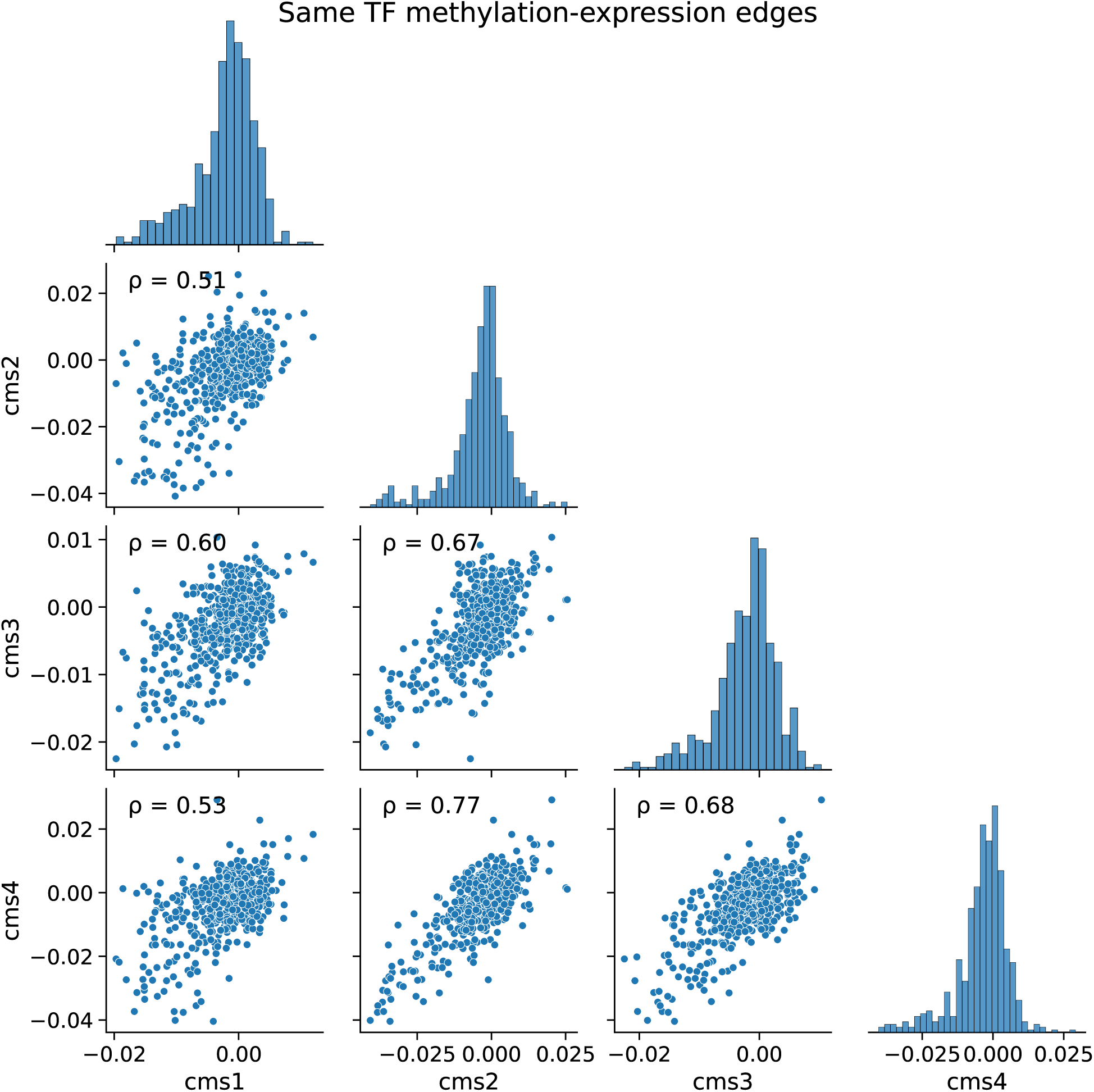
Correlation between DRAGON methylation-expression edges on the the same TFs. For all (*M_i_*, *E_i_*) edges we plot their distribution in each subtype (histograms on the diagonal) and the correlation of the edge weights between each pair of subtypes. While all Pearson correlation values are above 0.50 it is worth noting that CMS2 and CMS4 are the most similar to each other with ρ = 0.77. This shows that many of the TFs for which we have evidence of negative partial correlations between methylation and expression are conserved across the CMS2 and CMS4 subtypes, while being more different for CMS1.

**Figure S6.**
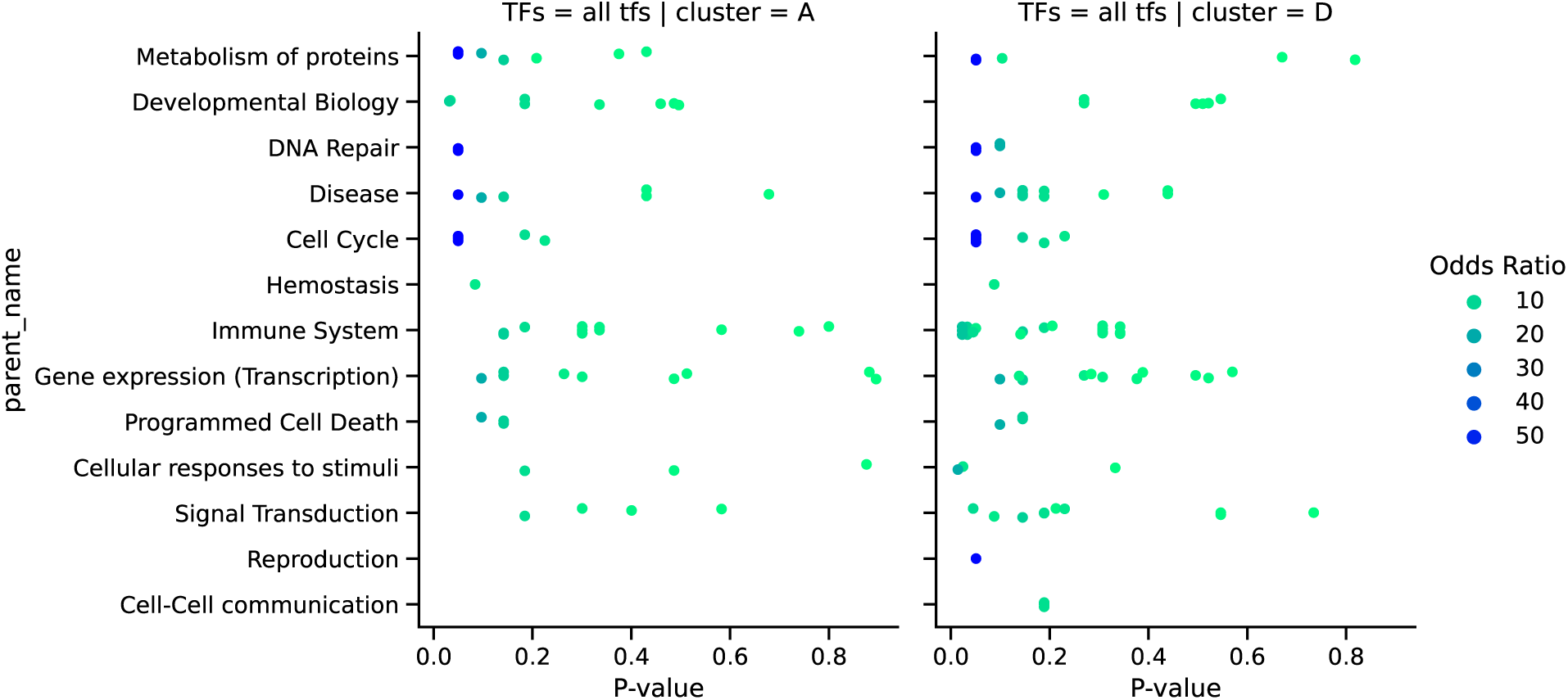
Reactome pathways for clusters A and D. Using the clusters found from the DRAGON edges, we run a pathway overrepresentation analysis of the TFs in both cluster A and D. With REACTOME we are able to identify the general pathway to which each term belongs to. For each “parent“” pathway (y-axis), we plot the P-values (x-axis) of all the pathways tested that belong to that parent term, and we color them by the corresponding Odds-Ratio. Since pathway analysis on TFs is challenging (there are only ∼ 1000 TFs and many of them are annotated only to the general transcriptional pathway terms) we report here all results, even those that are not significant, such that one can observe the general trend. TFs in cluster D seems to be more consistently annotated to Immune System pathways, while the TFs in cluster A have some stronger terms related to Developmental Biology.

**Figure S7.**
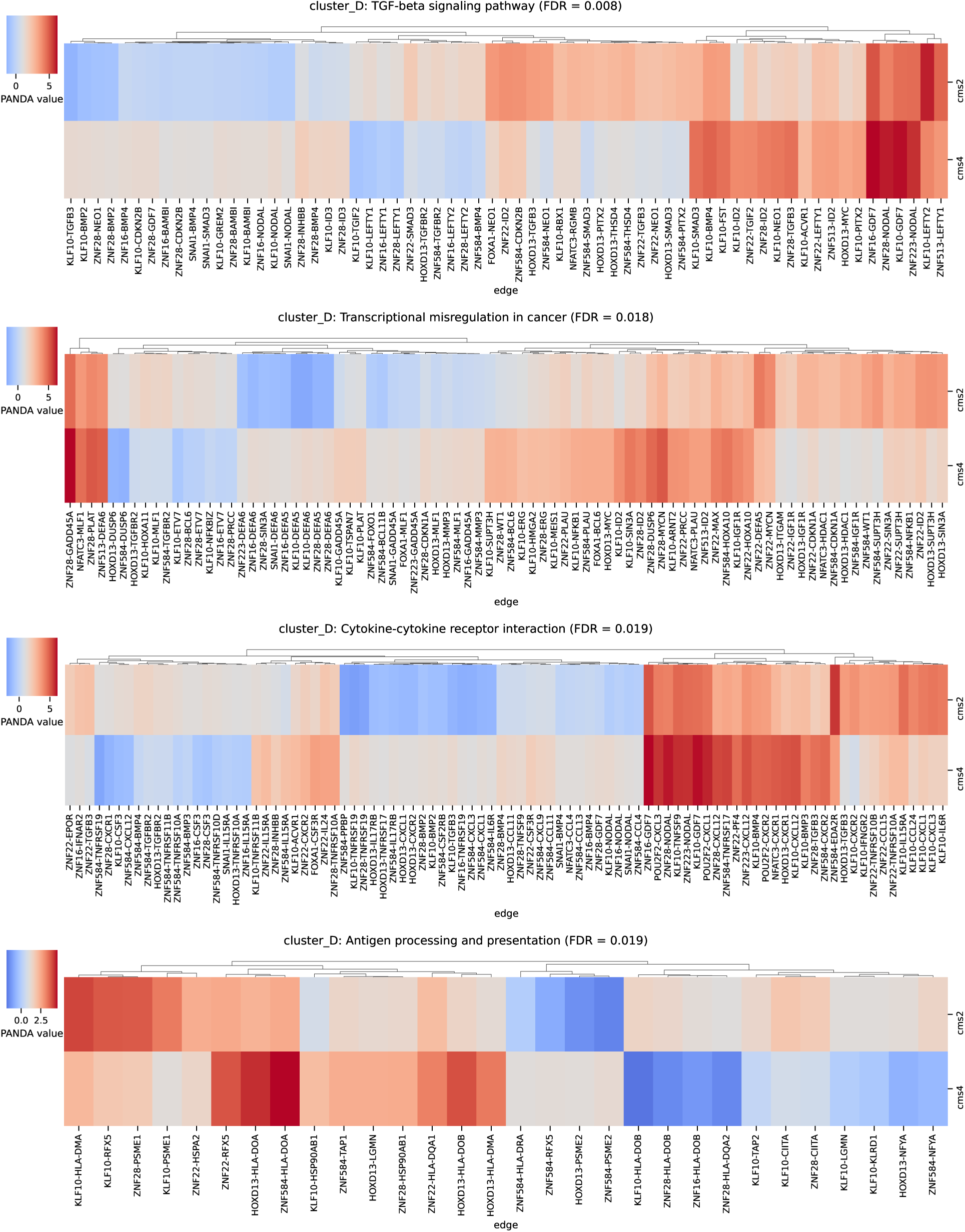
PANDA edges involved in the main pahtways targeted by cluster D. We selected the regulatory edges of the TFs in the cluster D (defined by the analysis on DRAGON networks) and we investigated which edges in the PANDA networks underwent the biggest changes between CMS2 and CMS4. The target genes of the edges were found to be preferentially involved in the TGF-beta signaling pathway, Transcriptional misregulation in cancer, Cytokine-cytokine receptor interaction and Antigen processing anr presentation. Here, we represent the PANDA edges (edge weight represented by different colors) connecting the main targets for each pathway, for both CMS2 and CMS4 (rows).

## Supplementary Tables

**Table S1.**
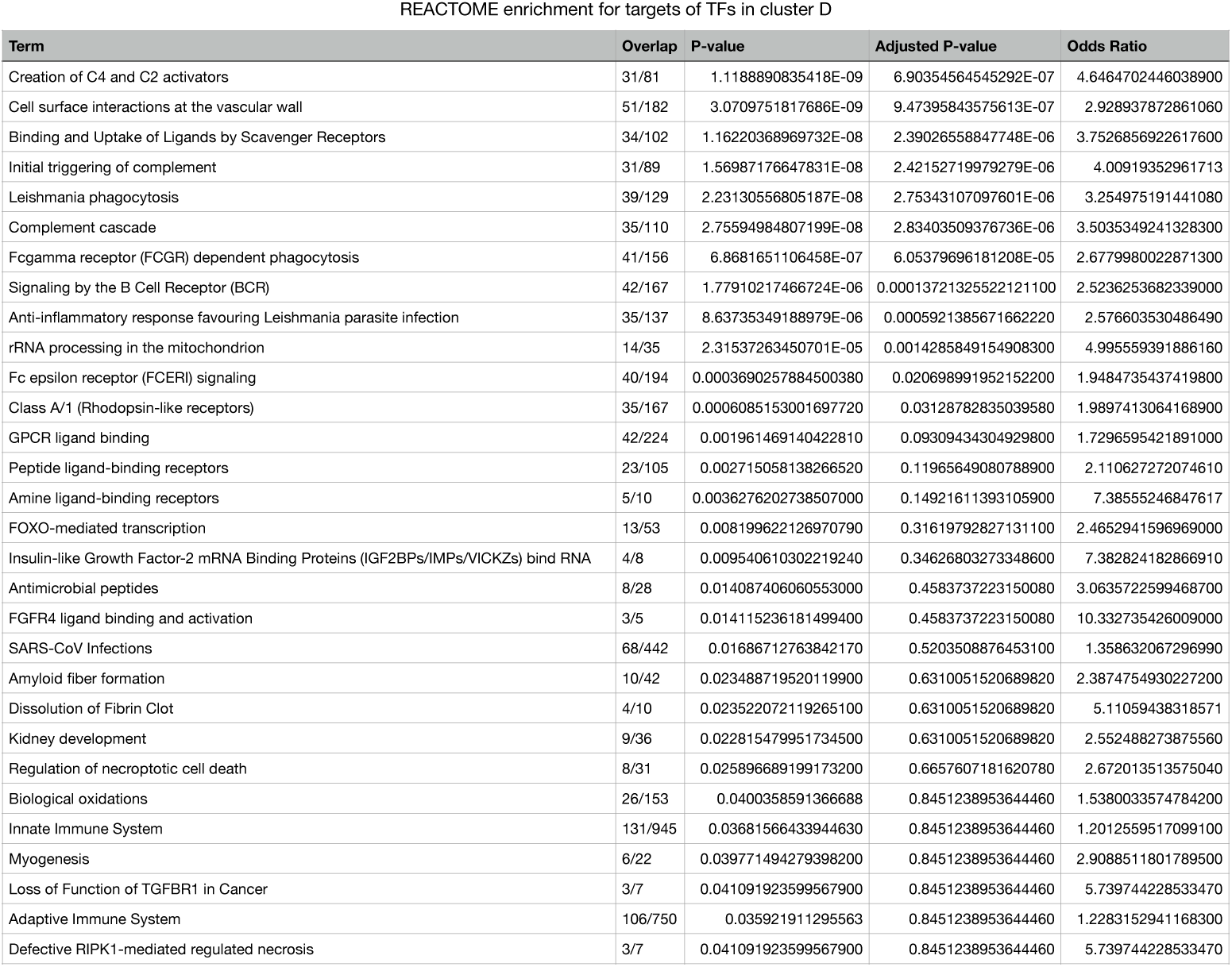
REACTOME pathway enrichment for the targets of TFs in cluster D. For the targets of the TFs in cluster D, we run a pathway overrepresentation analysis with the REACTOME pathway database. Here we show the pathways with *p* – *value* < 0.05.

**Table S2.**
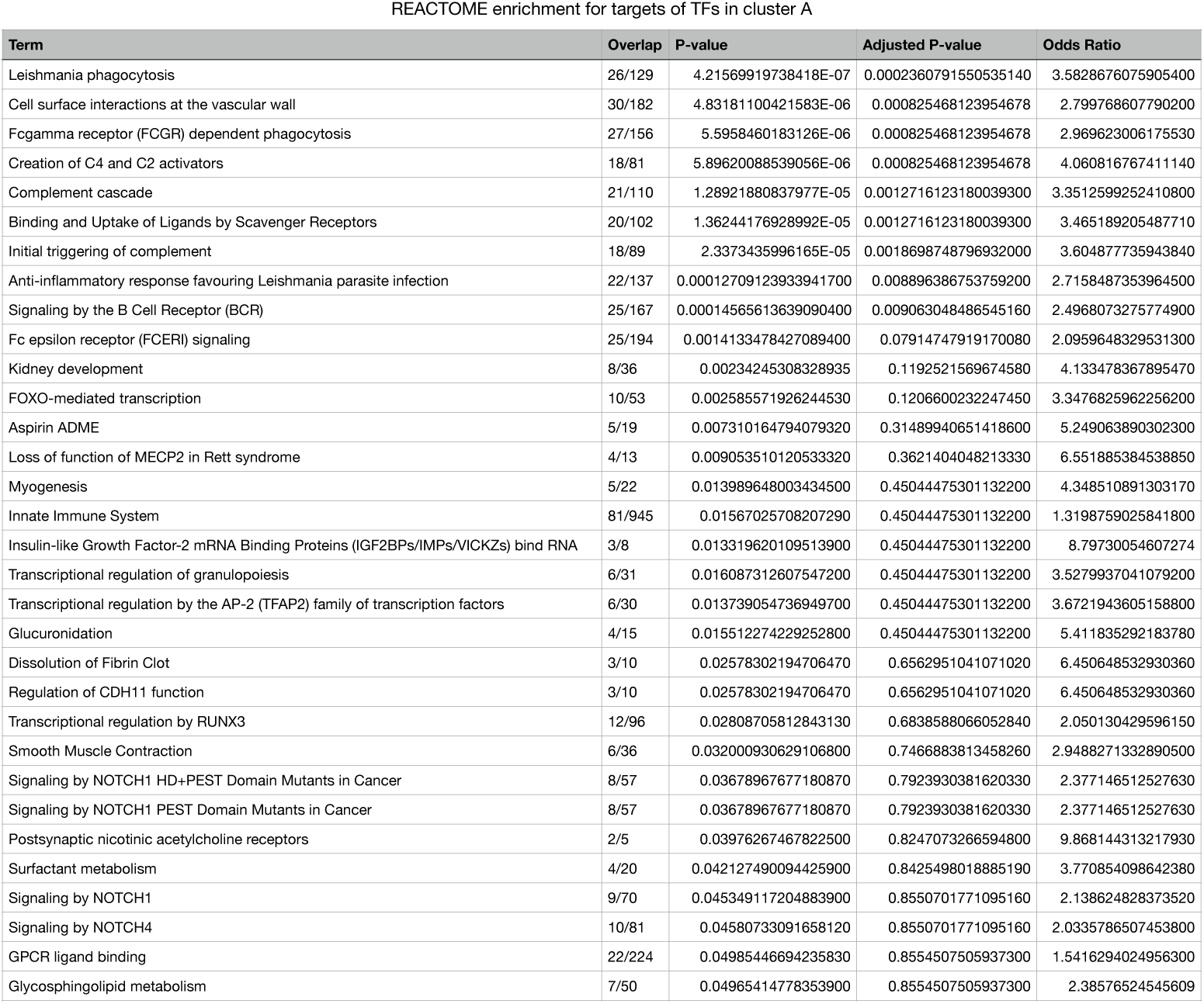
REACTOME pathway enrichment for the targets of TFs in cluster A. For the targets of the TFs in cluster A, we run a pathway overrepresentation analysis with the REACTOME pathway database. Here we show the pathways with *p* – *value* < 0.05.

**Table S3.** KEGG pathway enrichment for the targets of TFs in cluster D. For the targets of the TFs in cluster D, we run a pathway overrepresentation analysis with the KEGG pathway database. Here we show the pathways with *p* – *value* < 0.05.

| Term | P-value | Adjusted P-value | Odds Ratio |
| --- | --- | --- | --- |
| TGF-beta signaling pathway | 2.48574311940069E-05 | 0.0077058036701421300 | 3.019988310929280 |
| Transcriptional misregulation in cancer | 0.00011838600511097900 | 0.018349830792201800 | 2.2167008430166300 |
| Cytokine-cytokine receptor interaction | 0.00018606258744939800 | 0.018861201326119900 | 2.01001783033033 |
| Antigen processing and presentation | 0.00024337033969187000 | 0.018861201326119900 | 3.1492372152986700 |
| Amoebiasis | 0.003167301213570950 | 0.16063641285107300 | 2.2336917562724000 |
| Asthma | 0.003300331201939580 | 0.16063641285107300 | 4.703154782744 |
| Viral protein interaction with cytokine and cytokine receptor | 0.003627273838572620 | 0.16063641285107300 | 2.25914860748794 |
| Gastric cancer | 0.004165072298866050 | 0.16139655158106000 | 1.930047973264340 |
| Pathways in cancer | 0.005309881763878660 | 0.18289592742248700 | 1.4278343896029800 |
| Hematopoietic cell lineage | 0.00642522348971196 | 0.19918192818107100 | 2.169620497014060 |
| Epstein-Barr virus infection | 0.00857343648688083 | 0.22511457232477200 | 1.6532418398493700 |
| Arachidonic acid metabolism | 0.008714112477087950 | 0.22511457232477200 | 2.6239508869930700 |
| Neuroactive ligand-receptor interaction | 0.011684597675844700 | 0.2734075242836080 | 1.7469509466507200 |
| Calcium signaling pathway | 0.013035464353594000 | 0.2734075242836080 | 1.6528891842498800 |
| Signaling pathways regulating pluripotency of stem cells | 0.013803833745416300 | 0.2734075242836080 | 1.7899688597363000 |
| Retinol metabolism | 0.014111356092057200 | 0.2734075242836080 | 2.54912389563573 |
| Staphylococcus aureus infection | 0.01520219063915510 | 0.27721641753753300 | 2.185045435045440 |
| IL-17 signaling pathway | 0.01630080366536720 | 0.2807360631257680 | 1.972852233676980 |
| Inflammatory bowel disease | 0.020345197764012200 | 0.33194796351809400 | 2.2588584083440500 |
| Intestinal immune network for IgA production | 0.02281522942217870 | 0.3536360560437700 | 2.4633832976445400 |
| Primary immunodeficiency | 0.02589642462614270 | 0.36567840481762300 | 2.5699821322215600 |
| Linoleic acid metabolism | 0.025951370664476400 | 0.36567840481762300 | 3.6923899102180400 |
| Complement and coagulation cascades | 0.029464306184411400 | 0.3971276050942400 | 1.9615125329411000 |
| Type I diabetes mellitus | 0.031024169351406000 | 0.40072885412232700 | 2.4627568493150700 |
| Autoimmune thyroid disease | 0.03533995207094060 | 0.4382154056796640 | 2.585387248609330 |
| Graft-versus-host disease | 0.042424200647072800 | 0.5032938481671670 | 2.4621309370988400 |
| Prostate cancer | 0.043835270646817800 | 0.5032938481671670 | 1.6761839278040400 |
| Hepatocellular carcinoma | 0.04938104058133400 | 0.5467186635790540 | 1.4917963805311000 |

**Table S4.** KEGG pathway enrichment for the targets of TFs in cluster A. For the targets of the TFs in cluster A, we run a pathway overrepresentation analysis with the KEGG pathway database. Here we show the pathways with *p* – *value* < 0.05.

| Term | P-value | Adjusted P-value | Odds Ratio |
| --- | --- | --- | --- |
| Pathways in cancer | 0.000747938 | 0.14230677574040300 | 1.7114527709087400 |
| Transcriptional misregulation in cancer | 0.000978053 | 0.14230677574040300 | 2.302796633179470 |
| Gastric cancer | 0.003165216 | 0.250779096 | 2.272352647352650 |
| Intestinal immune network for IgA production | 0.009353948 | 0.250779096 | 3.3438080050194800 |
| Hippo signaling pathway | 0.005438134 | 0.250779096 | 2.0849770642201800 |
| Hepatocellular carcinoma | 0.005589136 | 0.250779096 | 2.031952209993530 |
| Basal cell carcinoma | 0.007511519 | 0.250779096 | 2.9016263177411700 |
| Cytokine-cytokine receptor interaction | 0.004160565 | 0.250779096 | 1.9401753529151800 |
| PPAR signaling pathway | 0.015161895289182000 | 0.250779096 | 2.545488354795220 |
| Endometrial cancer | 0.013600153655384300 | 0.250779096 | 2.5986617312072900 |
| Bacterial invasion of epithelial cells | 0.012675416382725400 | 0.250779096 | 2.475711029092490 |
| Bile secretion | 0.012160736019803900 | 0.250779096 | 2.6540978044879600 |
| Hematopoietic cell lineage | 0.011645194336257000 | 0.250779096 | 2.383638307984790 |
| Colorectal cancer | 0.011599440441574000 | 0.250779096 | 2.280352786639140 |
| Epstein-Barr virus infection | 0.010780171259324300 | 0.250779096 | 1.8245258620689700 |
| TGF-beta signaling pathway | 0.00965665 | 0.250779096 | 2.3448453276738100 |
| Fatty acid biosynthesis | 0.015512109 | 0.250779096 | 5.030944849401730 |
| Linoleic acid metabolism | 0.015512109 | 0.250779096 | 5.030944849401730 |
| Metabolism of xenobiotics by cytochrome P450 | 0.018044423 | 0.2723088279033320 | 2.6387744779247100 |
| Adherens junction | 0.018715383361053700 | 0.2723088279033320 | 2.310157042 |
| Porphyrin and chlorophyll metabolism | 0.021584665678258800 | 0.28550626 | 3.0764309764309800 |
| Steroid hormone biosynthesis | 0.021584665678258800 | 0.28550626 | 3.0764309764309800 |
| Mineral absorption | 0.026930818315737400 | 0.3407333969512860 | 2.61967502 |
| Thyroid cancer | 0.028226189 | 0.34224254437779000 | 2.863949843260190 |
| Asthma | 0.029600377927220100 | 0.3445483990728420 | 3.952236870542470 |
| Viral protein interaction with cytokine and cytokine receptor | 0.033977179 | 0.3802830416208250 | 2.068003448 |
| Drug metabolism | 0.036708966396366000 | 0.395641082 | 2.037479886 |
| Ether lipid metabolism | 0.040496699562009600 | 0.4208764133051710 | 2.5950284090909100 |
| Retinol metabolism | 0.045230707223669600 | 0.45386675179613300 | 2.5162534435261700 |
| Ascorbate and aldarate metabolism | 0.049349391 | 0.47868908895841100 | 3.254249354810000 |

## Notes

https://doi.org/10.7910/DVN/MCSSYJ

https://tcga-data-nf-precomputed.s3.us-east-2.amazonaws.com/raw-data/firstround-20221102/

https://github.com/QuackenbushLab/NetworkDataCompanion/

https://github.com/QuackenbushLab/tcga-data-nf

https://github.com/QuackenbushLab/tcga-data-supplement

### Competing Interest Statement

The authors have declared no competing interest.

## References

1. Mesirov JP. Accessible Reproducible Research. Science 2010 Jan;327(5964):415–416.

2. Baker M. 1,500 Scientists Lift the Lid on Reproducibility. Nature 2016 May;533(7604):452–454.

3. Heil BJ, Hoffman MM, Markowetz F, Lee SI, Greene CS, Hicks SC. Reproducibility Standards for Machine Learning in the Life Sciences. Nature methods 2021 Oct;18(10):1132–1135.

4. Munafò MR, Nosek BA, Bishop DVM, Button KS, Chambers CD, Percie du Sert N, et al. A Manifesto for Reproducible Science. Nature Human Behaviour 2017 Jan;1(1):1–9.

5. Gentleman RC, Carey VJ, Bates DM, Bolstad B, Dettling M, Dudoit S, et al. Bioconductor: Open Software Development for Computational Biology and Bioinformatics. Genome Biology 2004;.

6. Grüning B, Dale R, Sjödin A, Chapman BA, Rowe J, Tomkins-Tinch CH, et al. Bioconda: Sustainable and Comprehensive Software Distribution for the Life Sciences. Nature Methods 2018 Jul;15(7):475–476.

7. Jalili V, Afgan E, Gu Q, Clements D, Blankenberg D, Goecks J, et al. The Galaxy Platform for Accessible, Reproducible and Collaborative Biomedical Analyses: 2020 Update. Nucleic Acids Research 2020 Jul;48(W1):W395–W402.

8. Di Tommaso P, Chatzou M, Floden EW, Barja PP, Palumbo E, Notredame C. Nextflow Enables Reproducible Computational Workflows. Nature Biotechnology 2017 Apr;35(4):316–319.

9. Köster J, Rahmann S. Snakemake—a Scalable Bioinformatics Workflow Engine. Bioinformatics 2012 Oct;28(19):2520–2522.

10. Voss K, der Auwera GV, Gentry J. Full-stack Genomics Pipelining with GATK4 + WDL + Cromwell. F1000Research 2017 Aug;6.

11. Sudlow C, Gallacher J, Allen N, Beral V, Burton P, Danesh J, et al. UK Biobank: An Open Access Resource for Identifying the Causes of a Wide Range of Complex Diseases of Middle and Old Age. PLOS Medicine 2015 Mar;12(3):e1001779.

12. Fairley S, Lowy-Gallego E, Perry E, Flicek P. The International Genome Sample Resource (IGSR) Collection of Open Human Genomic Variation Resources. Nucleic Acids Research 2020 Jan;48(D1):D941–D947.

13. Turnbull C, Scott RH, Thomas E, Jones L, Murugaesu N, Pretty FB, et al. The 100 000 Genomes Project: bringing whole genome sequencing to the NHS. BMJ 2018 Apr;361:k1687.

14. Weinstein JN, Collisson EA, Mills GB, Shaw KRM, Ozenberger BA, Ellrott K, et al. The Cancer Genome Atlas Pan-Cancer Analysis Project. Nature Genetics 2013 Oct;45(10):1113–1120.

15. Bailey MH, Tokheim C, Porta-Pardo E, Sengupta S, Bertrand D, Weerasinghe A, et al. Comprehensive Characterization of Cancer Driver Genes and Mutations. Cell 2018 Apr;173(2):371–385.e18.

16. Sanchez-Vega F, Mina M, Armenia J, Chatila WK, Luna A, La KC, et al. Oncogenic Signaling Pathways in The Cancer Genome Atlas. Cell 2018 Apr;173(2):321–337.e10.

17. Ding L, Bailey MH, Porta-Pardo E, Thorsson V, Colaprico A, Bertrand D, et al. Perspective on Oncogenic Processes at the End of the Beginning of Cancer Genomics. Cell 2018 Apr;173(2):305–320.e10.

18. Hoadley KA, Yau C, Hinoue T, Wolf DM, Lazar AJ, Drill E, et al. Cell-of-Origin Patterns Dominate the Molecular Classification of 10,000 Tumors from 33 Types of Cancer. Cell 2018 Apr;173(2):291–304.e6.

19. Weighill D, Ben Guebila M, Glass K, Platig J, Yeh JJ, Quackenbush J. Gene targeting in disease networks. Front Genet 2021 Apr;12:649942.

20. Ellrott K, Bailey MH, Saksena G, Covington KR, Kandoth C, Stewart C, et al. Scalable Open Science Approach for Mutation Calling of Tumor Exomes Using Multiple Genomic Pipelines. Cell Systems 2018 Mar;6(3):271–281.e7.

21. Way GP, Sanchez-Vega F, La K, Armenia J, Chatila WK, Luna A, et al. Machine Learning Detects Pan-cancer Ras Pathway Activation in The Cancer Genome Atlas. Cell Reports 2018 Apr;23(1):172–180.e3.

22. Li J, Lu Y, Akbani R, Ju Z, Roebuck PL, Liu W, et al. TCPA: A Resource for Cancer Functional Proteomics Data. Nature Methods 2013 Nov;10(11):1046–1047.

23. Li J, Akbani R, Zhao W, Lu Y, Weinstein JN, Mills GB, et al. Explore, Visualize, and Analyze Functional Cancer Proteomic Data Using the Cancer Proteome Atlas. Cancer Research 2017 Nov;77(21):e51–e54.

24. Edwards NJ, Oberti M, Thangudu RR, Cai S, McGarvey PB, Jacob S, et al. The CPTAC Data Portal: A Resource for Cancer Proteomics Research. Journal of Proteome Research 2015 Jun;14(6):2707–2713.

25. Ellis MJ, Gillette M, Carr SA, Paulovich AG, Smith RD, Rodland KK, et al. Connecting Genomic Alterations to Cancer Biology with Proteomics: The NCI Clinical Proteomic Tumor Analysis Consortium. Cancer Discovery 2013 Oct;3(10):1108–1112.

26. Cowen L, Ideker T, Raphael BJ, Sharan R. Network Propagation: A Universal Amplifier of Genetic Associations. Nature Reviews Genetics 2017 Sep;18(9):551–562.

27. Sonawane AR, Platig J, Fagny M, Chen CY, Paulson JN, Lopes-Ramos CM, et al. Understanding Tissue-Specific Gene Regulation. Cell reports 2017;21(4):1077–1088.

28. Reyna MA, Haan D, Paczkowska M, Verbeke LPC, Vazquez M, Kahraman A, et al. Pathway and Network Analysis of More than 2500 Whole Cancer Genomes. Nature Communications 2020 Feb;11(1):729.

29. Leiserson MDM, Vandin F, Wu HT, Dobson JR, Eldridge JV, Thomas JL, et al. Pan-Cancer Network Analysis Identifies Combinations of Rare Somatic Mutations across Pathways and Protein Complexes. Nature Genetics 2015 Feb;47(2):106–114.

30. Silverbush D, Cristea S, Yanovich-Arad G, Geiger T, Beerenwinkel N, Sharan R. Simultaneous Integration of Multi-omics Data Improves the Identification of Cancer Driver Modules. Cell Systems 2019 May;8(5):456–466.e5.

31. Belova T, Biondi N, Hsieh PH, Lutsik P, Chudasama P, Kuijjer ML. Heterogeneity in the Gene Regulatory Landscape of Leiomyosarcoma. NAR Cancer 2023 Sep;5(3):zcad037.

32. Lopes-Ramos CM, Kuijjer ML, Ogino S, Fuchs CS, DeMeo DL, Glass K, et al. Gene regulatory network analysis identifies sex-linked differences in colon cancer drug metabolism. Cancer Res 2018 Oct;78(19):5538–5547.

33. Glass K, Huttenhower C, Quackenbush J, Yuan GC. Passing Messages between Biological Networks to Refine Predicted Interactions. PLOS ONE 2013 May;8(5):e64832.

34. Weighill D, Guebila MB, Lopes-Ramos C, Glass K, Quackenbush J, Platig J, et al. Gene Regulatory Network Inference as Relaxed Graph Matching. Proceedings of the AAAI Conference on Artificial Intelligence 2021 May;35(11):10263–10272.

35. Chen C, Padi M, Joint Inference of Transcription Factor Activity and Context-Specific Regulatory Networks. bioRxiv; 2022.

36. Margolin AA, Nemenman I, Basso K, Wiggins C, Stolovitzky G, Favera RD, et al. ARACNE: An Algorithm for the Reconstruction of Gene Regulatory Networks in a Mammalian Cellular Context. BMC Bioinformatics 2006 Mar;7(1):S7.

37. Alvarez MJ, Shen Y, Giorgi FM, Lachmann A, Ding BB, Ye BH, et al. Network-Based Inference of Protein Activity Helps Functionalize the Genetic Landscape of Cancer. Nature genetics 2016 Aug;48(8):838–847.

38. Saha E, Ben Guebila M, Fanfani V, Fischer J, Shutta KH, Mandros P, et al. Gene regulatory networks reveal sex difference in lung adenocarcinoma. Biol Sex Differ 2024 Aug;15(1):62.

39. Shutta KH, Weighill D, Burkholz R, Guebila MB, DeMeo DL, Zacharias HU, et al. DRAGON: Determining Regulatory Associations Using Graphical Models on Multi-Omic Networks. Nucleic Acids Research 2023 Feb;51(3):e15.

40. GenomicDataCommons;. http://bioconductor.org/packages/GenomicDataCommons/.

41. Morgan MT, Davis SR, GenomicDataCommons: A Bioconductor Interface to the NCI Genomic Data Commons. bioRxiv; 2017.

42. Colaprico A, Silva TC, Olsen C, Garofano L, Cava C, Garolini D, et al. TCGAbiolinks: An R/Bioconductor Package for Integrative Analysis of TCGA Data. Nucleic Acids Research 2016 May;44(8):e71.

43. Mounir M, Lucchetta M, Silva TC, Olsen C, Bontempi G, Chen X, et al. New Functionalities in the TCGAbiolinks Package for the Study and Integration of Cancer Data from GDC and GTEx. PLoS computational biology 2019 Mar;15(3):e1006701.

44. Silva TC, Colaprico A, Olsen C, D’Angelo F, Bontempi G, Ceccarelli M, et al., TCGA Workflow: Analyze Cancer Genomics and Epigenomics Data Using Bioconductor Packages; 2016.

45. Ben Guebila M, Wang T, Lopes-Ramos CM, Fanfani V, Weighill D, Burkholz R, et al. The Network Zoo: A Multilingual Package for the Inference and Analysis of Gene Regulatory Networks. Genome Biology 2023 Mar;24(1):45.

46. Kuijjer ML, Tung MG, Yuan G, Quackenbush J, Glass K. Estimating Sample-Specific Regulatory Networks. iScience 2019 Apr;14:226– 240.

47. Guinney J, Dienstmann R, Wang X, de Reyniès A, Schlicker A, Soneson C, et al. The Consensus Molecular Subtypes of Colorectal Cancer. Nature Medicine 2015 Nov;21(11):1350–1356.

48. Ben Guebila M, Lopes-Ramos CM, Weighill D, Sonawane AR, Burkholz R, Shamsaei B, et al. GRAND: A Database of Gene Regulatory Network Models across Human Conditions. Nucleic Acids Research 2022 Jan;50(D1):D610–D621.

49. Merkel D. Docker: Lightweight Linux Containers for Consistent Development and Deployment. Linux journal 2014;2014(239):2.

50. Kurtzer GM, Sochat V, Bauer MW. Singularity: Scientific Containers for Mobility of Compute. PLOS ONE 2017 May;12(5):e0177459.

51. Anaconda Software Distribution. Anaconda Inc.; 2020.

52. Grossman Robert L, Heath Allison P, Ferretti Vincent, Varmus Harold E, Lowy Douglas R, Kibbe Warren A, et al. Toward a Shared Vision for Cancer Genomic Data. New England Journal of Medicine 2016;375(12):1109–1112.

53. Wilks C, Zheng SC, Chen FY, Charles R, Solomon B, Ling JP, et al. Recount3: Summaries and Queries for Large-Scale RNA-seq Expression and Splicing. Genome biology 2021;.

54. THE GTEX CONSORTIUM. The GTEx Consortium Atlas of Genetic Regulatory Effects across Human Tissues. Science 2020 Sep;369(6509):1318–1330.

55. Arora S, Pattwell SS, Holland EC, Bolouri H, Uncertainty in RNA-seq Gene Expression Data; 2018.

56. Johnson KA, Krishnan A. Robust Normalization and Transformation Techniques for Constructing Gene Coexpression Networks from RNA-seq Data. Genome Biology 2022 Jan;23(1):1.

57. Collado-Torres L, Nellore A, Kammers K, Ellis SE, Taub MA, Hansen KD, et al. Reproducible RNA-seq Analysis Using Recount2. Nature Biotechnology 2017 Apr;35(4):319–321.

58. Li B, Ruotti V, Stewart RM, Thomson JA, Dewey CN. RNA-Seq Gene Expression Estimation with Read Mapping Uncertainty. Bioinformatics 2010 Feb;26(4):493–500.

59. Robinson MD, McCarthy DJ, Smyth GK. edgeR: A Bioconductor Package for Differential Expression Analysis of Digital Gene Expression Data. Bioinformatics 2010 Jan;26(1):139–140.

60. Chen Y, Chen L, Lun ATL, Baldoni PL, Smyth GK, edgeR 4.0: Powerful Differential Analysis of Sequencing Data with Expanded Functionality and Improved Support for Small Counts and Larger Datasets. bioRxiv; 2024.

61. Johnson WE, Li C, Rabinovic A. Adjusting Batch Effects in Microarray Expression Data Using Empirical Bayes Methods. Biostatistics (Oxford, England) 2007 Jan;8(1):118–127.

62. Leek JT, Johnson WE, Parker HS, Jaffe AE, Storey JD. The Sva Package for Removing Batch Effects and Other Unwanted Variation in High-Throughput Experiments. Bioinformatics 2012 Mar;28(6):882–883.

63. Aran D, Sirota M, Butte AJ. Systematic Pan-Cancer Analysis of Tumour Purity. Nature Communications 2015 Dec;6(1):8971.

64. Du P, Zhang X, Huang CC, Jafari N, Kibbe WA, Hou L, et al. Comparison of Beta-value and M-value Methods for Quantifying Methylation Levels by Microarray Analysis. BMC bioinformatics 2010;11:1–9.

65. Liu H, Lafferty J, Wasserman L. The Nonparanormal: Semiparametric Estimation of High Dimensional Undirected Graphs. Journal of Machine Learning Research 2009;10(10).

66. Zhao T, Liu H, Roeder K, Lafferty J, Wasserman L. The Huge Package for High-Dimensional Undirected Graph Estimation in R. The Journal of Machine Learning Research 2012;13(1):1059–1062.

67. Martin FJ, Amode MR, Aneja A, Austine-Orimoloye O, Azov AG, Barnes I, et al. Ensembl 2023. Nucleic Acids Research 2023 Jan;51(D1):D933–D941.

68. Seal RL, Braschi B, Gray K, Jones TEM, Tweedie S, Haim-Vilmovsky L, et al. Genenames.Org: The HGNC Resources in 2023. Nucleic Acids Research 2023 Jan;51(D1):D1003–D1009.

69. Brown GR, Hem V, Katz KS, Ovetsky M, Wallin C, Ermolaeva O, et al. Gene: A Gene-Centered Information Resource at NCBI. Nucleic Acids Research 2015 Jan;43(Database issue):D36–42.

70. Maglott D, Ostell J, Pruitt KD, Tatusova T. Entrez Gene: Gene-Centered Information at NCBI. Nucleic Acids Research 2011 Jan;39(suppl_1):D52–D57.

71. Frankish A, Diekhans M, Jungreis I, Lagarde J, Loveland JE, Mudge JM, et al. GENCODE 2021. Nucleic Acids Research 2021 Jan;49(D1):D916–D923.

72. AnnotationDbi;. http://bioconductor.org/packages/AnnotationDbi/.

73. Du P, Zhang X, Huang CC, Jafari N, Kibbe WA, Hou L, et al. Comparison of Beta-value and M-value Methods for Quantifying Methylation Levels by Microarray Analysis. BMC Bioinformatics 2010 Nov;11(1):587.

74. Ferlay J, Ervik M, Lam F, Colombet M, Mery L, Piñeros M, et al. Global Cancer Observatory: Cancer Today. Lyon, France: international agency for research on cancer 2024;Available from: https://gco.iarc.who.int/today, accessed [19 June 2024].(0):0.

75. Marisa L, Blum Y, Taieb J, Ayadi M, Pilati C, Le Malicot K, et al. Intratumor CMS Heterogeneity Impacts Patient Prognosis in Localized Colon Cancer. Clinical Cancer Research 2021 Sep;27(17):4768–4780.

76. Mouillet-Richard S, Cazelles A, Sroussi M, Gallois C, Taieb J, Laurent-Puig P. Clinical Challenges of Consensus Molecular Subtype CMS4 Colon Cancer in the Era of Precision Medicine. Clinical Cancer Research 2024 Apr;p. OF1–OF8.

77. OncoKB: A Precision Oncology Knowledge Base | JCO Precision Oncology;. https://ascopubs.org/doi/full/10.1200/PO.17.00011.

78. Mattei AL, Bailly N, Meissner A. DNA Methylation: A Historical Perspective. Trends in Genetics 2022 Jul;38(7):676–707.

79. Chakravarty D, Gao J, Phillips S, Kundra R, Zhang H, Wang J, et al. OncoKB: A Precision Oncology Knowledge Base. JCO Precision Oncology 2017 Dec;(1):1–16.

80. Suehnholz SP, Nissan MH, Zhang H, Kundra R, Nandakumar S, Lu C, et al. Quantifying the Expanding Landscape of Clinical Actionability for Patients with Cancer. Cancer Discovery 2024 Jan;14(1):49–65.

81. Tan SH, Nevalainen MT. Signal Transducer and Activator of Transcription 5A/B in Prostate and Breast Cancers. Endocrine-Related Cancer 2008 Jun;15(2):367–390.

82. Haddad BR, Gu L, Mirtti T, Dagvadorj A, Vogiatzi P, Hoang DT, et al. STAT5A/B Gene Locus Undergoes Amplification during Human Prostate Cancer Progression. The American Journal of Pathology 2013 Jun;182(6):2264–2275.

83. Di Palma T, Lucci V, de Cristofaro T, Filippone MG, Zannini M. A Role for PAX8 in the Tumorigenic Phenotype of Ovarian Cancer Cells. BMC cancer 2014 Apr;14:292.

84. Jesse S, Koenig A, Ellenrieder V, Menke A. Lef-1 Isoforms Regulate Different Target Genes and Reduce Cellular Adhesion. International Journal of Cancer 2010 Mar;126(5):1109–1120.

85. Chiang YT, Wang K, Fazli L, Qi RZ, Gleave ME, Collins CC, et al. GATA2 as a Potential Metastasis-Driving Gene in Prostate Cancer. Oncotarget 2014 Jan;5(2):451–461.

86. Mellor P, Deibert L, Calvert B, Bonham K, Carlsen SA, Anderson DH. CREB3L1 Is a Metastasis Suppressor That Represses Expression of Genes Regulating Metastasis, Invasion, and Angiogenesis. Molecular and Cellular Biology 2013 Dec;33(24):4985–4995.

87. Heide T, Househam J, Cresswell GD, Spiteri I, Lynn C, Mossner M, et al. The Co-Evolution of the Genome and Epigenome in Colorectal Cancer. Nature 2022 Nov;611(7937):733–743.

88. Sondka Z, Bamford S, Cole CG, Ward SA, Dunham I, Forbes SA. The COSMIC Cancer Gene Census: Describing Genetic Dysfunction across All Human Cancers. Nature Reviews Cancer 2018 Nov;18(11):696–705.

89. Johnson KA, Krishnan A. Robust Normalization and Transformation Techniques for Constructing Gene Coexpression Networks from RNA-seq Data. Genome Biology 2022 Jan;23(1):1.

90. Haider S, Tyekucheva S, Prandi D, Fox NS, Ahn J, Xu AW, et al. Systematic Assessment of Tumor Purity and Its Clinical Implications. JCO Precision Oncology 2020 Nov;(4):995–1005.

91. Fang Z, Liu X, Peltz G. GSEApy: a comprehensive package for performing gene set enrichment analysis in Python. Bioinformatics 2022 11;39(1):btac757. 10.1093/bioinformatics/btac757.

92. Grant CE, Bailey TL, Noble WS. FIMO: scanning for occurrences of a given motif. Bioinformatics 2011;27(7):1017–1018.

93. Szklarczyk D, Gable AL, Nastou KC, Lyon D, Kirsch R, Pyysalo S, et al. The STRING database in 2021: customizable protein–protein networks, and functional characterization of user-uploaded gene/measurement sets. Nucleic acids research 2021;49(D1):D605–D612.

